# Large-Scale Comparative Genomics and Structure-Function Analysis Enables Characterization of Known and Novel Genetic Determinants of Antimicrobial Resistance in Bacterial Pathogens

**DOI:** 10.64898/2026.01.27.702097

**Authors:** Anthony Mannion, Daniel Hooks, Arianna Comendul, Rebecca Spirgel

## Abstract

Antibiotics are crucial for preventing infection-induced complications, but their widespread overuse has spurred the evolution of antimicrobial resistance (AMR) mechanisms in pathogens. Data-driven biosurveillance approaches utilizing whole genome sequencing data and computational approaches have the potential to improve the detection and characterization of known and emerging AMR profiles, especially in high-priority ESKAPE, enteric, and sexually-transmitted pathogens. In this study, we performed a large-scale analysis of over 70,000 genomes representing 39 pathogen-antibiotic combinations to identify resistance determinants statistically enriched in antibiotic resistant strains. Using a kmer-based GWAS approach, over 7,000 unique sequences were identified among all resistant genomes. Of these, 1,925 sequences were homologous to known AMR genes, while over 5,000 sequences lacked homology, suggesting novel AMR-associated genes. In addition to identifying the predominant AMR genes for specific pathogen-antibiotic combinations, our findings suggest that horizontal gene transfer mechanisms may influence AMR gene profiles between phylogenetically similar pathogens and antibiotic classes. Likewise, significant associations in co-harbored, multi-drug resistance mechanisms were identified in select pathogens. Protein domains analysis frequently detected efflux/membrane structure and antibiotic-associated metabolism domains in novel AMR-associated proteins, suggesting additional mechanisms potentiate resistance phenotypes. Furthermore, we developed a Random Forest classifier using protein structure, molecular features, and binding affinity profiles to predict protein-antibiotic interactions, identifying several novel proteins that may interact with antibiotics. This study demonstrates the potential of large-scale comparative genomics coupled with AI/ML-based modeling to advance the understanding of AMR threats, thereby enhancing biosurveillance efforts and promoting new strategies to counteract emerging pathogens.

## Introduction

Antibiotics are essential treatments in modern medicine due to their proven efficacy to reduce infection-induced complications from injury or other common procedures like surgery. Consequently, antibiotics are often administered prophylactically and frequently without prior diagnostic tests to confirm pathogen target susceptibility. Wide-spread overuse of antibiotics over the last century has enabled pathogens to acquire/evolve antimicrobial resistance (AMR) mechanisms to counteract these treatments. Currently, AMR pathogens are attributed to greater than 2.8 million infections and cause 35,000 deaths per year in the United States.^1^ Globally, over 5 million deaths annually are due to AMR infections.^1,2^ By 2050, AMR pathogens are predicted to cause greater than 10 million deaths per year, which will exceed those caused by cancer.^2^ Thus, AMR is a significant threat jeopardizing the treatment of pathogenic infections.

Over the last several decades, numerous resistance mechanisms have been identified in pathogens that drive AMR phenotypes.^3^ Identification of these mechanisms has relied on laboratory-based assays for culture/isolation of suspect resistant strains and screening for antibiotic susceptibility, followed by characterization and experimental confirmation of the genetic determinants.^4^ Thus, current approaches for AMR identification are labor-, cost-, and time-intensive.^4^ As a result of these significant challenges, understanding the prevalence and distribution of AMR genes, especially in high-priority pathogens such as ESKAPE+ (e.g., *Enterococcus faecium*, *Staphylococcus aureus*, *Klebsiella pneumoniae*, *Acinetobacter baumannii*, *Pseudomonas aeruginosa*, *Enterobacter species*, *Escherichia coli*), enteric and sexually-transmitted infections, is hindered.^5^ In turn, this has hampered the development of improved strategies to detect, prevent, and treat known and emerging AMR pathogens, such as novel diagnostic or treatment modalities.

Since the advent of next-generation sequence technologies, public databases that incorporate whole genome sequencing (WGS) with laboratory-based antibiograms have been developed for pathogenic strains and offer the potential to identify reservoirs of genes associated with AMR.^6^ Furthermore, recent advances in bioinformatic and computation biology capabilities via artificial intelligence (AI) and machine learning (ML) tools, such as 3D protein structure prediction and ligand docking modeling,^7^ provide opportunities to investigate mechanisms that promote to AMR, including potentially novel genetic determinants. Herein, we hypothesize that large-scale comparative genomic coupled with AI/ML-based structure-function modeling can be leveraged to enhance the identification and characterization of known and novel AMR mechanism in pathogens.

To this end, over 70,000 genomes, representing 39 unique pathogen-antibiotic combinations, were analyzed for AMR-associated proteins using a kmer-based profiling approach. AI/ML-based modeling was then used to characterize the mechanism of interaction between AMR proteins and the antibiotics molecules. In addition to profiling the frequency of known AMR gene variant per pathogen-antibiotic combination, over 5,000 gene sequences were identified that were statistically associated with drug-resistant phenotypes and lack sequence homology to known AMR genes in the Comprehensive Antibiotic Resistance Database (CARD).^8^ To augment traditional annotation tools such as protein domain analysis for functional inference, a novel ML classification algorithm was developed using structure-function modeling tools to predict interactions between AMR genes and antibiotic targets. The findings from this work provide new perspectives on the prevalence and distribution of known AMR gene variants and facilitate the mechanistic characterization of novel protein sequences associated with AMR using emerging AI/ML modeling approaches.

## Methods

### Open-Source Data and Comparative Genomics

The API for Bacterial and Viral Bioinformatics Resource Center (BV-BRC)^9^ was utilized to download annotated proteins from WGS for taxa with laboratory-based antibiograms present in the metadata (accessed March 25, 2025). Antibiotic susceptibility and resistance labels for genome were defined on the antibiogram metadata provided for the strain. Up to 1,500 random genomes for resistant and susceptible strains per pathogen-antibiotic combination were selected for analysis by DBGWAS (De Bruijn Graph Genome-wide association study), under default settings to identify kmers associated with AMR (**Table 1**).^10^ Kmers enriched resistant genomes with a Q-value of ≤1E-10 were then mapped to annotated protein sequences. DIAMOND (double index alignment of next-generation sequencing data) was performed against the Comprehensive Antibiotic Resistance Database (CARD, version 4.0.0) to identify homologs to known AMR genes and differentiate non-known AMR genes.^8,11^ Identical sequences for non-known AMR genes were de-duplicated to generate representative gene sets. InterProScan (version 5.64-96.0) was used to annotate protein domains for gene sequences.^12^ De-duplicated AMR genes were also clustered into representative gene sequenced using DIAMOND based on a percent identity of 30%.

**Table 1.**
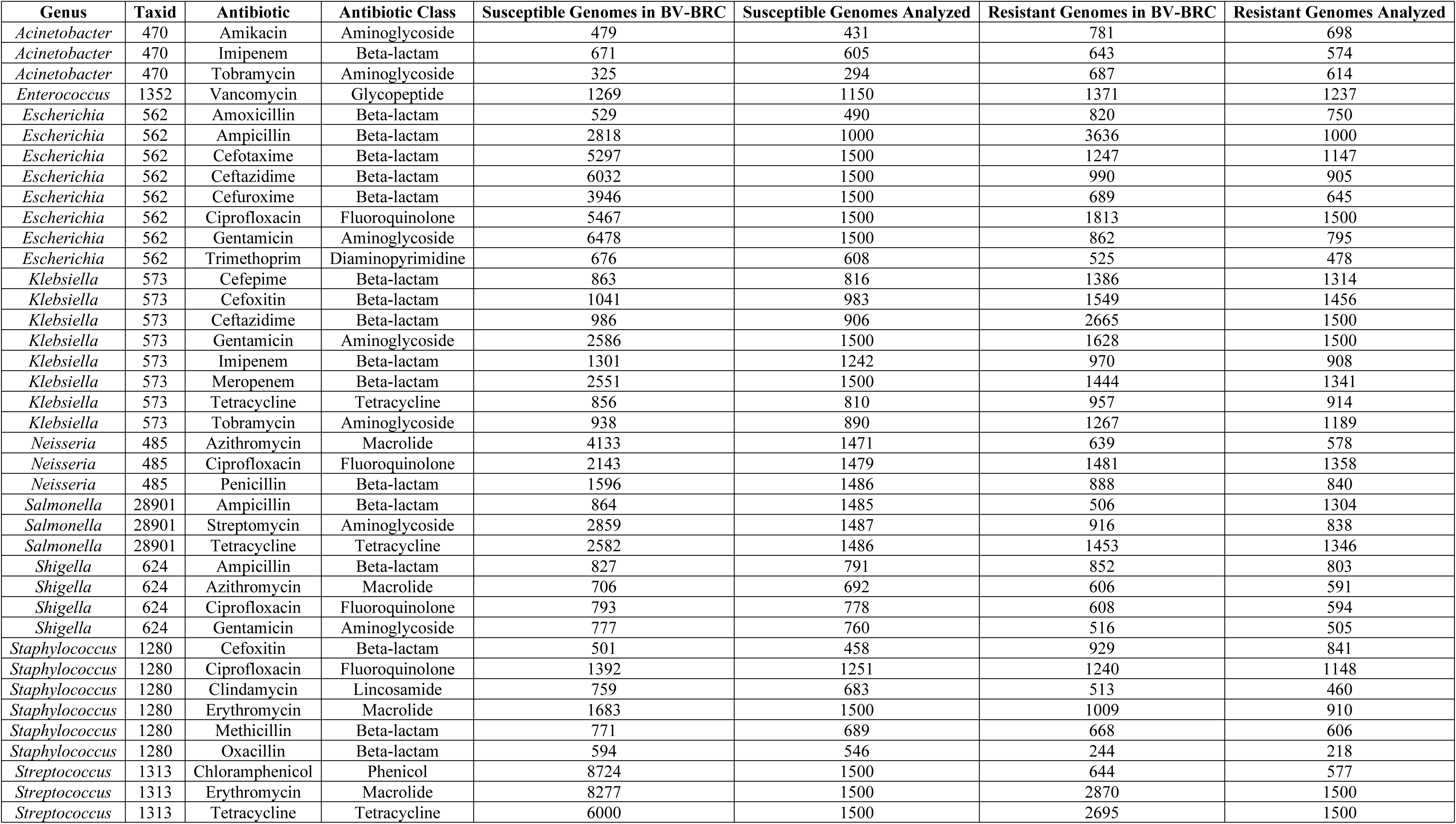
Summary of Genomes Analyzed.

| Genus | Taxid | Antibiotic | Antibiotic Class | Susceptible Genomes in BV-BRC | Susceptible Genomes Analyzed | Resistant Genomes in BV-BRC | Resistant Genomes Analyzed |
| --- | --- | --- | --- | --- | --- | --- | --- |
| *Acinetobacter* | 470 | Amikacin | Aminoglycoside | 479 | 431 | 781 | 698 |
| *Acinetobacter* | 470 | Imipenem | Beta-lactam | 671 | 605 | 643 | 574 |
| *Acinetobacter* | 470 | Tobramycin | Aminoglycoside | 325 | 294 | 687 | 614 |
| *Enterococcus* | 1352 | Vancomycin | Glycopeptide | 1269 | 1150 | 1371 | 1237 |
| *Escherichia* | 562 | Amoxicillin | Beta-lactam | 529 | 490 | 820 | 750 |
| *Escherichia* | 562 | Ampicillin | Beta-lactam | 2818 | 1000 | 3636 | 1000 |
| *Escherichia* | 562 | Cefotaxime | Beta-lactam | 5297 | 1500 | 1247 | 1147 |
| *Escherichia* | 562 | Ceftazidime | Beta-lactam | 6032 | 1500 | 990 | 905 |
| *Escherichia* | 562 | Cefuroxime | Beta-lactam | 3946 | 1500 | 689 | 645 |
| *Escherichia* | 562 | Ciprofloxacin | Fluoroquinolone | 5467 | 1500 | 1813 | 1500 |
| *Escherichia* | 562 | Gentamicin | Aminoglycoside | 6478 | 1500 | 862 | 795 |
| *Escherichia* | 562 | Trimethoprim | Diaminopyrimidine | 676 | 608 | 525 | 478 |
| *Klebsiella* | 573 | Cefepime | Beta-lactam | 863 | 816 | 1386 | 1314 |
| *Klebsiella* | 573 | Cefoxitin | Beta-lactam | 1041 | 983 | 1549 | 1456 |
| *Klebsiella* | 573 | Ceftazidime | Beta-lactam | 986 | 906 | 2665 | 1500 |
| *Klebsiella* | 573 | Gentamicin | Aminoglycoside | 2586 | 1500 | 1628 | 1500 |
| *Klebsiella* | 573 | Imipenem | Beta-lactam | 1301 | 1242 | 970 | 908 |
| *Klebsiella* | 573 | Meropenem | Beta-lactam | 2551 | 1500 | 1444 | 1341 |
| *Klebsiella* | 573 | Tetracycline | Tetracycline | 856 | 810 | 957 | 914 |
| *Klebsiella* | 573 | Tobramycin | Aminoglycoside | 938 | 890 | 1267 | 1189 |
| *Neisseria* | 485 | Azithromycin | Macrolide | 4133 | 1471 | 639 | 578 |
| *Neisseria* | 485 | Ciprofloxacin | Fluoroquinolone | 2143 | 1479 | 1481 | 1358 |
| *Neisseria* | 485 | Penicillin | Beta-lactam | 1596 | 1486 | 888 | 840 |
| *Salmonella* | 28901 | Ampicillin | Beta-lactam | 864 | 1485 | 506 | 1304 |
| *Salmonella* | 28901 | Streptomycin | Aminoglycoside | 2859 | 1487 | 916 | 838 |
| *Salmonella* | 28901 | Tetracycline | Tetracycline | 2582 | 1486 | 1453 | 1346 |
| *Shigella* | 624 | Ampicillin | Beta-lactam | 827 | 791 | 852 | 803 |
| *Shigella* | 624 | Azithromycin | Macrolide | 706 | 692 | 606 | 591 |
| *Shigella* | 624 | Ciprofloxacin | Fluoroquinolone | 793 | 778 | 608 | 594 |
| *Shigella* | 624 | Gentamicin | Aminoglycoside | 777 | 760 | 516 | 505 |
| *Staphylococcus* | 1280 | Cefoxitin | Beta-lactam | 501 | 458 | 929 | 841 |
| *Staphylococcus* | 1280 | Ciprofloxacin | Fluoroquinolone | 1392 | 1251 | 1240 | 1148 |
| *Staphylococcus* | 1280 | Clindamycin | Lincosamide | 759 | 683 | 513 | 460 |
| *Staphylococcus* | 1280 | Erythromycin | Macrolide | 1683 | 1500 | 1009 | 910 |
| *Staphylococcus* | 1280 | Methicillin | Beta-lactam | 771 | 689 | 668 | 606 |
| *Staphylococcus* | 1280 | Oxacillin | Beta-lactam | 594 | 546 | 244 | 218 |
| *Streptococcus* | 1313 | Chloramphenicol | Phenicol | 8724 | 1500 | 644 | 577 |
| *Streptococcus* | 1313 | Erythromycin | Macrolide | 8277 | 1500 | 2870 | 1500 |
| *Streptococcus* | 1313 | Tetracycline | Tetracycline | 6000 | 1500 | 2695 | 1500 |

### 3D Protein Structure and Ligand Docking

ESM-Fold (version 2) was used to generate 3D protein structure.^13^ DiffDock2 was then used to dock a seed ligand (e.g., isoniazid) in 3D protein structure to identify the generalized ligand binding region.^14^ Next, GNINA (version 1.1) under the vinardo scoring function was used to refine the ligand pose within the binding box and calculate the predicted binding affinity.^15^

### ML Models for Classification of Protein-Ligand Interactions

Data was collected from the CARD database consisting of 6,300 proteins and 641 ligands, resulting 4,038,300 protein-ligand combinations. For every combination that had a known interaction, it was labeled as 1. For all other combinations, those without a known interaction, a label of 0 was applied. Only proteins that confer resistance as inactivation enzymes to beta-lactam, aminoglycoside, tetracycline, or macrolides were used for training and testing due to class imbalances for other mechanism of action types and molecule types. Randomly selected subsets of proteins and ligands were used for the test set. Any combination that included a protein or a ligand in the test set was excluded from training, according to the following conditions: Given the set of Ligands, *L* = (*l*_1_, *l*_2_, … , *l*_*n*_)where *n* = |*L*|, and a set of Proteins, *P* = (*p*_1_, *p*_2_, … , *p*_*m*_)where *m* = |*P*|. We randomly sampled from *L* and from *P* to create the set *L*^′^, *P*^′^. The training set was created by [(*L* − *L*^′^) × (*P* − *P*^′^)] − (*L*, *P* ∈ [*d*(*l*, *l*_*i*_) × *b*(*p*, *p*_*i*_), *l*_*i*_ × *p*_*i*_ = 1, *l* × *p* ≠ 1])where *d*(*x*, *y*)is the Tanimoto similarity of *l* and *l*_*i*_that is greater than a threshold *t* , and *b*(*x*, *y*) is the sequence similarity (based on DIAMOND analysis) between protein *p* and *p*_*i*_ >30%. The test set is the cross of the randomly sampled ligand and protein: *L*^′^ × *P*^′^.

ESM-Fold was used to represent the proteins as embedded vectors. After determining ligand docking sites with DiffDock2, GNINA was used to calculate the Affinity (kcal/mol), root mean squared error, convolutional neural network score, convolutional neural network affinity, and convolutional neural network variance. For each ligand, using their SMILES string representation, RDKit was used to calculate circular fingerprints, MACCSkey fingerprint, Avalon Fingerprint, ERG fingerprint, and rdkit features. Protein embedding, affinity output, and molecular features for each combination were used as features and the label of known or unknown interaction as labels. Using the training set, Random Forest and XGboost classifier models were trained. The probability thresholds for each classifier model were assessed to optimize sensitivity versus precision such that the known combinations were correctly classified without overestimating the unknown. A probability threshold of 0.5 was selected for the Random Forest model and 0.25 for the XGboost model to balance tradeoffs between sensitivity and precision for identification of known versus unknown protein-ligand interactions (**Supplemental Figure 2**). At these thresholds, performance metrics (precision, sensitivity, specificity, AUC, accuracy, PRAUC) were compared between the models across the average of all interaction pairs and per antibiotic class.

### Statistical Analysis

Chi-square tests were performed to analyze the statistical co-association AMR genes. A p-value <0.01 was considered statistically significant.

### Computing Resources and Code Availability

For DBGWAS, an Intel Xeon Platinum 8260 Processor with 375 Gb of RAM was used. For DiffDock and GNINA, up to two Nvidia Volta V100 GPUs were used. For training and testing ML classifiers, Intel Xeon E5-2683 v3 Processors with 28 cores and 256 Gb of RAM used. All code and models are available at upon request.

## Results

### DBGWAS-Based Identification of AMR-Associated AMR Genes

136,297 total genomes from nine different genera, representing 11 unique taxonomies, with laboratory-based antibiogram data for 24 different antibiotics were assessed from the BV-BRC database (**Table 1**). In total, 39 unique pathogen-antibiotic combinations were evaluated. Due to computational/memory limitations using DBGWAS, a maximum of 1,500 genomes from resistant and susceptible genomes (n = 3,000 total genomes maximum) could be compared per pathogen-antibiotic combination (**Table 1**), which represented 79,249 total genomes.

Across the 39 pathogen-antibiotic combinations, 7,149 unique sequences were identified that were enriched in the genomes of resistant strains. Of these sequences, 1,925 (29.9%) unique were homologous to known AMR-associated genes in the CARD, whereas 5,224 (73.1%) lacked homology hits (henceforth referred as “novel” AMR-associated genes). The distribution of known versus “novel” AMR-associated genes varied considerably per pathogen-antibiotic combination (**Figure 1**). Novel AMR-associated genes contributed >50% of total enriched sequences for 31/39 pathogen-antibiotic combinations.

**Figure 1:**
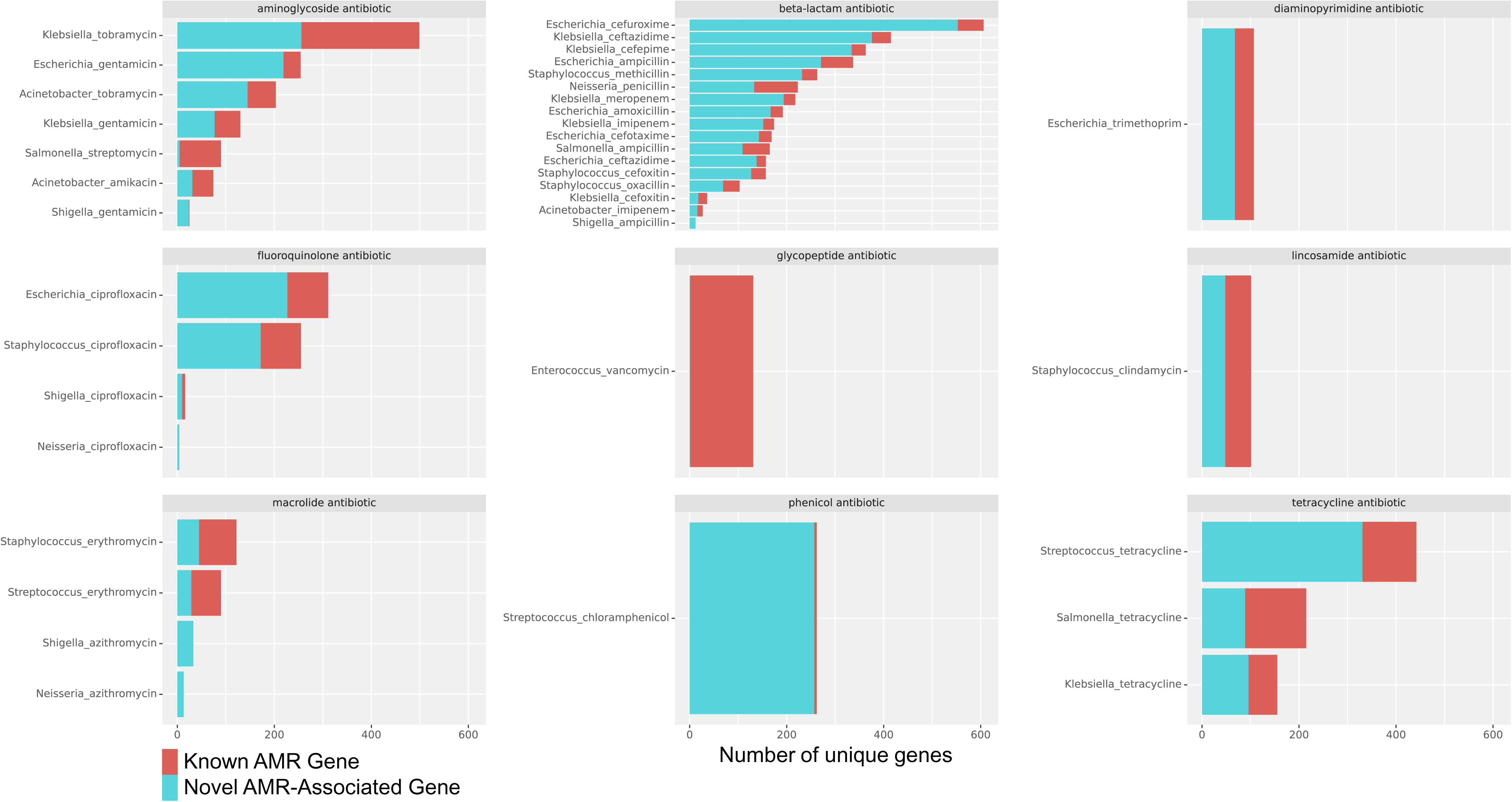
Total number of unique sequences detected for known and novel AMR-associated genes.

Of the 39 per pathogen-antibiotic combinations, 35 had known AMR-associated genes detected. The four pathogen-antibiotic combinations that lacked homology to known AMR-associated genes were *Shigella*-azithromycin, *Neisseria*-azithromycin, *Shigella*-ampicillin, and *Neisseria*-ciprofloxacin. Two pathogen-antibiotic combinations, *Enterococcus*-vancomycin and *Salmonella*-streptomycin, primarily harbored known AMR genes, with less <5% genomes co-harboring novel AMR-associated sequences. For the remaining 37 pathogen-antibiotic combos, novel AMR-associated sequences were identified in >10% of the genomes analyzed. In general, a higher percentage of genomes per pathogen-antibiotic combination tended to co-harbor both known and novel AMR-associated sequences. There were subsets of genomes that lacked both known versus “novel” AMR-associated genes (**Figure 2**), suggesting the presence of undetected AMR-associated sequences that were may have been statistically underpowered for detection using DBGWAS, possibly due low strain abundance in the BV-BRC database.

**Figure 2:**
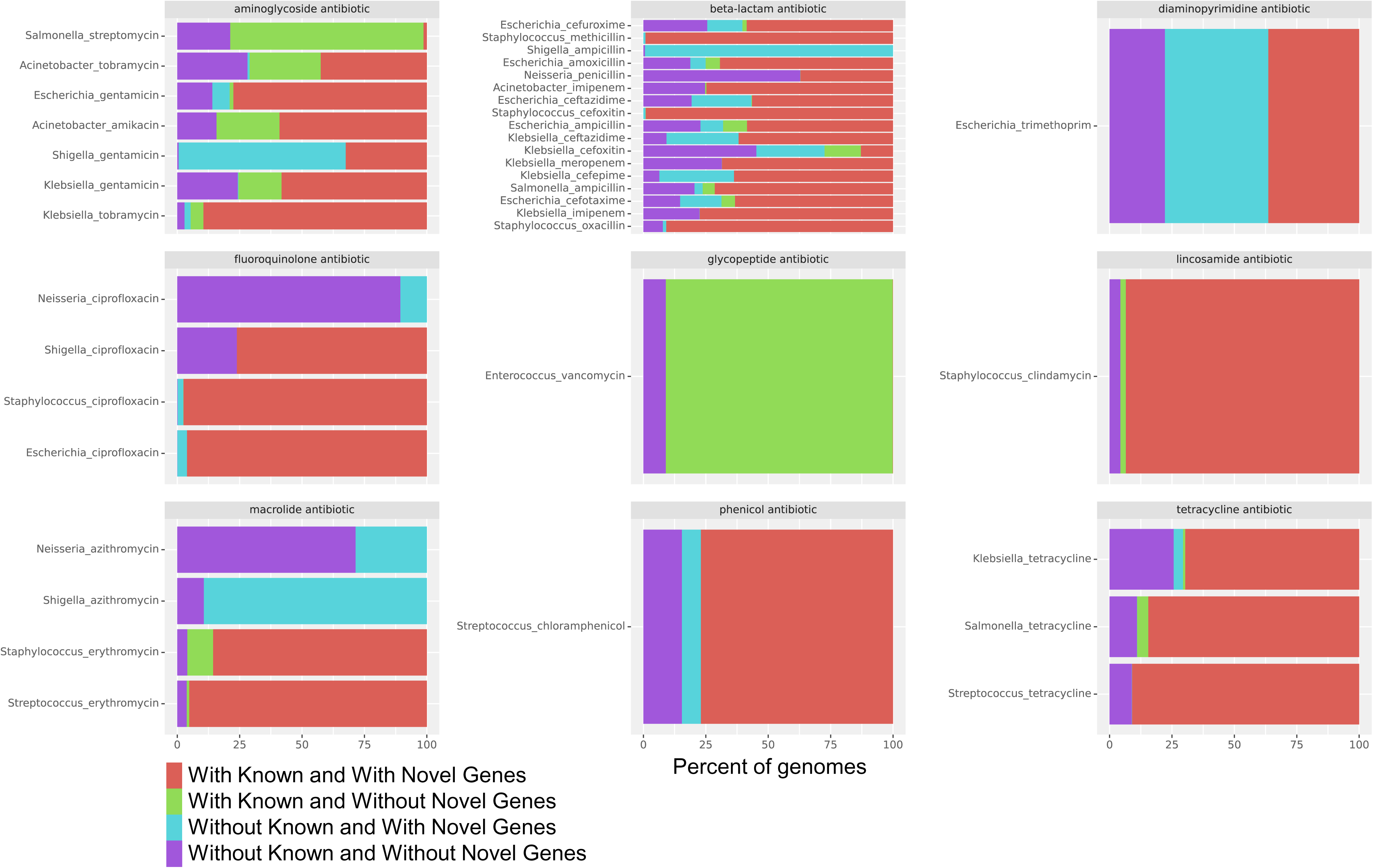
Percent of genomes with known and novel genes.

### On-Target AMR Gene Profiles

Known AMR genes were then assessed for whether they conferred resistance to the expected antibiotic class in pathogen–antibiotic combinations (i.e., distinguishing on-target from off-target AMR genes). For 35/39 pathogen-antibiotic combos, at least one on-target AMR gene detected. However, most pathogen-antibiotic combos had several on-target AMR genes identified, with up to 25 unique gene sequences for beta-lactam resistance (**Figure 3**).

**Figure 3:**
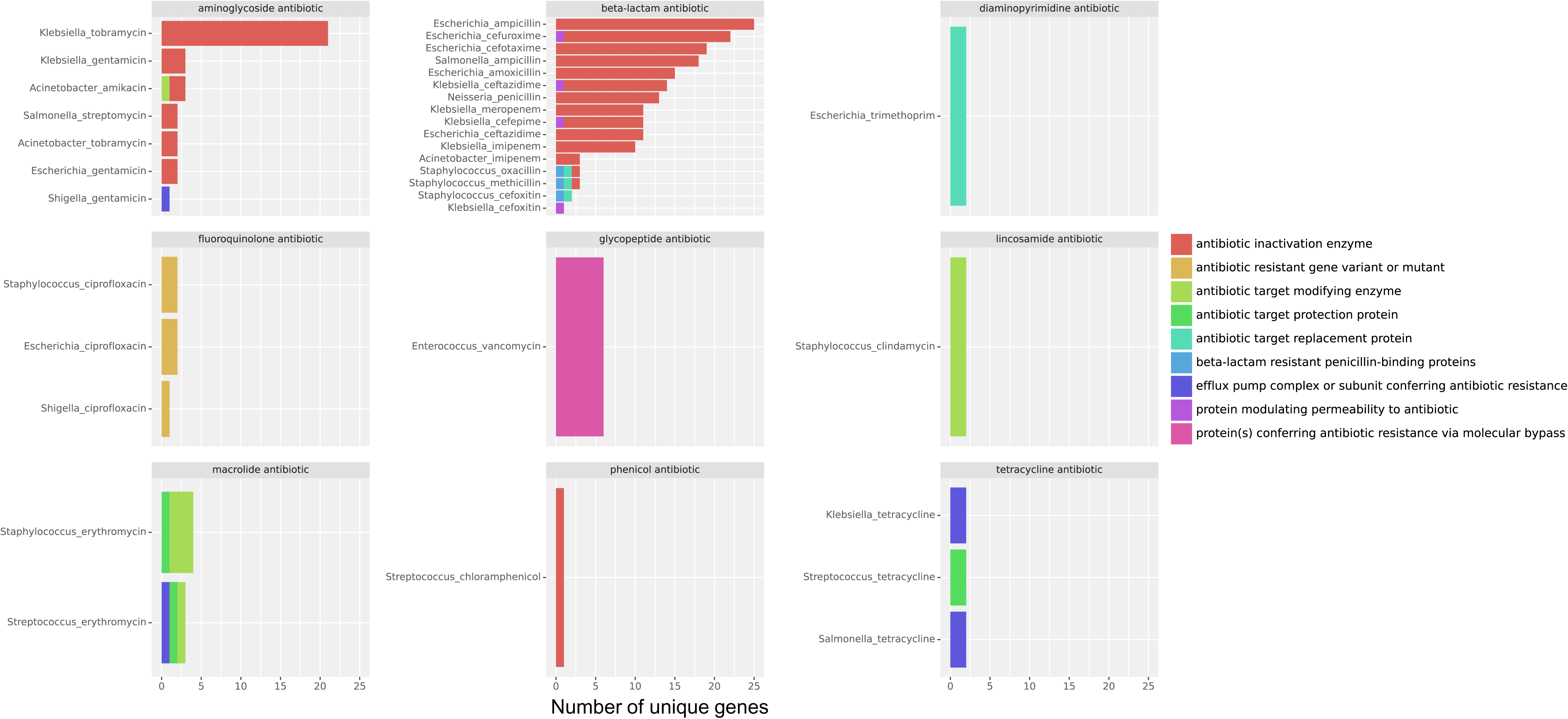
Number of unique on-target AMR genes and mechanism of action.

To determine if genomes tended to co-harbor specific on-target AMR gene pairs, chi-square tests were performed. Among all pathogen-genome combinations, only 17 on-target AMR gene pairs were found to be significantly associated when analyzing pairs found in >5% of genome (p-value <0.01, odds-ratio >1) (**Table 2**). The majority of co-harbored AMR genes (10/17 pairs) were found in *Enterococcus* genomes and were due to the gene cluster that confers resistance to vancomycin (**Table 2**). Interestingly, the remaining co-harbored AMR genes represented the same mechanisms of resistance, with the exception of a single pair found in *Streptococcus*-erythromycin in which the two genes had different mechanism of action. This observation suggests that it may be uncommon for genomes to harbor multiple different genes targeting same antibiotic, possibly because some resistance mechanism introduce redundancy that may reduce the overall fitness of the organism. It should be noted that efflux pumps were not considered in the co-associated gene comparison because these mechanisms potentiate non-specific resistance to broad classes of antibiotics.

**Table 2.**
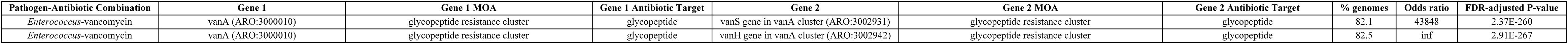
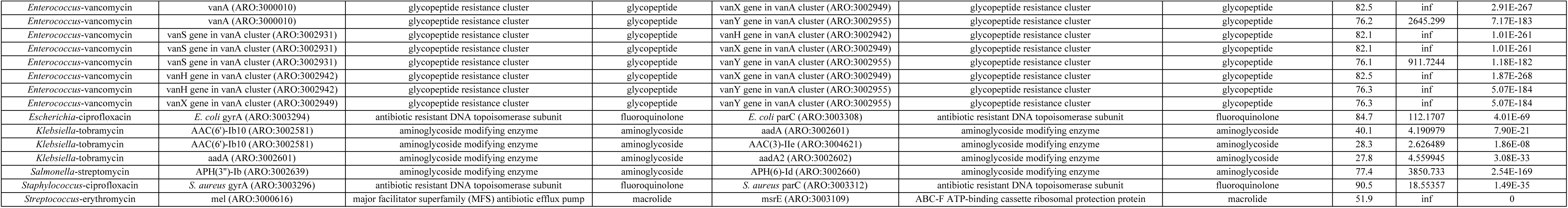
Characteristics of Statistically Co-Association of On-Target AMR Gene Pairs.

| Pathogen-Antibiotic Combination | Gene 1 | Gene 1 MOA | Gene 1 Antibiotic Target | Gene 2 | Gene 2 MOA | Gene 2 Antibiotic Target | % genomes | Odds ratio | FDR-adjusted P-value |
| --- | --- | --- | --- | --- | --- | --- | --- | --- | --- |
| *Enterococcus*-vancomycin | vanA (ARO:3000010) | glycopeptide resistance cluster | glycopeptide | vanS gene in vanA cluster (ARO:3002931) | glycopeptide resistance cluster | glycopeptide | 82.1 | 43848 | 2.37E-260 |
| *Enterococcus*-vancomycin | vanA (ARO:3000010) | glycopeptide resistance cluster | glycopeptide | vanH gene in vanA cluster (ARO:3002942) | glycopeptide resistance cluster | glycopeptide | 82.5 | inf | 2.91E-267 |
| *Enterococcus-vancomycin* | vanA (ARO:3000010) | glycopeptide resistance cluster | glycopeptide | vanX gene in vanA cluster (ARO:3002949) | glycopeptide resistance cluster | glycopeptide | 82.5 | inf | 2.91E-267 |
| *Enterococcus-vancomycin* | vanA (ARO:3000010) | glycopeptide resistance cluster | glycopeptide | vanY gene in vanA cluster (ARO:3002955) | glycopeptide resistance cluster | glycopeptide | 76.2 | 2645.299 | 7.17E-183 |
| *Enterococcus-vancomycin* | vanS gene in vanA cluster (ARO:3002931) | glycopeptide resistance cluster | glycopeptide | vanH gene in vanA cluster (ARO:3002942) | glycopeptide resistance cluster | glycopeptide | 82.1 | inf | 1.01E-261 |
| *Enterococcus-vancomycin* | vanS gene in vanA cluster (ARO:3002931) | glycopeptide resistance cluster | glycopeptide | vanX gene in vanA cluster (ARO:3002949) | glycopeptide resistance cluster | glycopeptide | 82.1 | inf | 1.01E-261 |
| *Enterococcus-vancomycin* | vanS gene in vanA cluster (ARO:3002931) | glycopeptide resistance cluster | glycopeptide | vanY gene in vanA cluster (ARO:3002955) | glycopeptide resistance cluster | glycopeptide | 76.1 | 911.7244 | 1.18E-182 |
| *Enterococcus-vancomycin* | vanH gene in vanA cluster (ARO:3002942) | glycopeptide resistance cluster | glycopeptide | vanX gene in vanA cluster (ARO:3002949) | glycopeptide resistance cluster | glycopeptide | 82.5 | inf | 1.87E-268 |
| *Enterococcus-vancomycin* | vanH gene in vanA cluster (ARO:3002942) | glycopeptide resistance cluster | glycopeptide | vanY gene in vanA cluster (ARO:3002955) | glycopeptide resistance cluster | glycopeptide | 76.3 | inf | 5.07E-184 |
| *Enterococcus-vancomycin* | vanX gene in vanA cluster (ARO:3002949) | glycopeptide resistance cluster | glycopeptide | vanY gene in vanA cluster (ARO:3002955) | glycopeptide resistance cluster | glycopeptide | 76.3 | inf | 5.07E-184 |
| *Escherichia-ciprofloxacin* | *E. coli* gyrA (ARO:3003294) | antibiotic resistant DNA topoisomerase subunit | fluoroquinolone | *E. coli* parC (ARO:3003308) | antibiotic resistant DNA topoisomerase subunit | fluoroquinolone | 84.7 | 112.1707 | 4.01E-69 |
| *Klebsiella-tobramycin* | AAC(6′)-Ib10 (ARO:3002581) | aminoglycoside modifying enzyme | aminoglycoside | aadA (ARO:3002601) | aminoglycoside modifying enzyme | aminoglycoside | 40.1 | 4.190979 | 7.90E-21 |
| *Klebsiella-tobramycin* | AAC(6′)-Ib10 (ARO:3002581) | aminoglycoside modifying enzyme | aminoglycoside | AAC(3)-Ile (ARO:3004621) | aminoglycoside modifying enzyme | aminoglycoside | 28.3 | 2.626489 | 1.86E-08 |
| *Klebsiella-tobramycin* | aadA (ARO:3002601) | aminoglycoside modifying enzyme | aminoglycoside | aadA2 (ARO:3002602) | aminoglycoside modifying enzyme | aminoglycoside | 27.8 | 4.559945 | 3.08E-33 |
| *Salmonella-streptomycin* | APH(3′)-Ib (ARO:3002639) | aminoglycoside modifying enzyme | aminoglycoside | APH(6)-Id (ARO:3002660) | aminoglycoside modifying enzyme | aminoglycoside | 77.4 | 3850.733 | 2.54E-169 |
| *Staphylococcus-ciprofloxacin* | *S. aureus* gyrA (ARO:3003296) | antibiotic resistant DNA topoisomerase subunit | fluoroquinolone | *S. aureus* parC (ARO:3003312) | antibiotic resistant DNA topoisomerase subunit | fluoroquinolone | 90.5 | 18.55357 | 1.49E-35 |
| *Streptococcus-erythromycin* | mel (ARO:3000616) | major facilitator superfamily (MFS) antibiotic efflux pump | macrolide | msrE (ARO:3003109) | ABC-F ATP-binding cassette ribosomal protection protein | macrolide | 51.9 | inf | 0 |

In general, on-target AMR genes tended to converge on a predominate mechanism for resistance based on the antibiotic class analyzed. For example, AMR genes functioning as antibiotic inactivation enzymes were the most commonly occurring mechanisms for beta-lactam and aminoglycoside classes for all pathogen-antibiotic combination (**Figure 3**). Likewise, specific AMR genes were predominantly identified for each pathogen-antibiotic combination. Additionally, hierarchal clustering revealed that pathogen-antibiotic combinations with more similar phylogeny and/or antibiotic classes tended to have more related AMR gene profiles (**Figure 4**). For example, *Escherichia*, *Klebsiella*, and *Salmonella* genomes (all of members of the *Enterobacteriaceae* family) resistant to penicillin-like molecule were enriched in TEM-1 (ARO:3008730). Conversely, *Escherichia* and *Klebsiella* genomes resistant to carbapenem or cephalosporin were enriched in KPC-2 beta-lactamase (ARO:3002312) / KPC-3 (ARO:3002313) or CTX-M-25 (ARO:3001887), respectively. Furthermore, *Acinetobacter* genomes, which are phylogenetically distant from *Enterobacteriaceae* species, that were resistant to carbapenems primarily harbored OXA-23 (ARO:3001418). A similar trend was also observed when novel AMR-associated genes were analyzed by hierarchal clustering in that gene profiles tended to more similar for pathogen-antibiotic combinations based on phylogeny and/or antibiotic class similarities (**Figure 5**, **Supplemental Figure 1**).

**Figure 4A.**
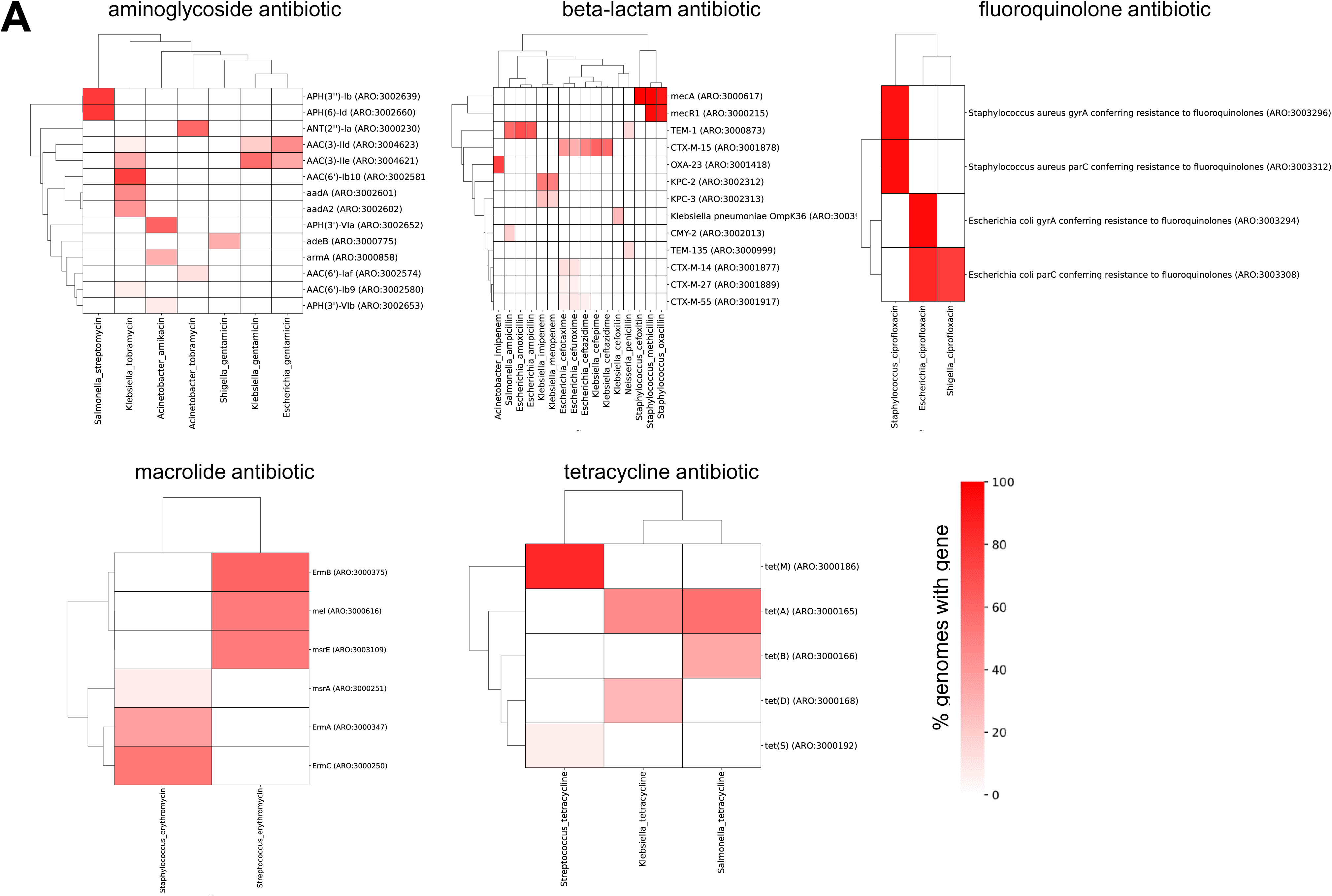
Clustermaps (by Euclidean distance) of known AMR gene abundance. Only genes present in ≥5% of genomes are shown.

**Figure 4B.**
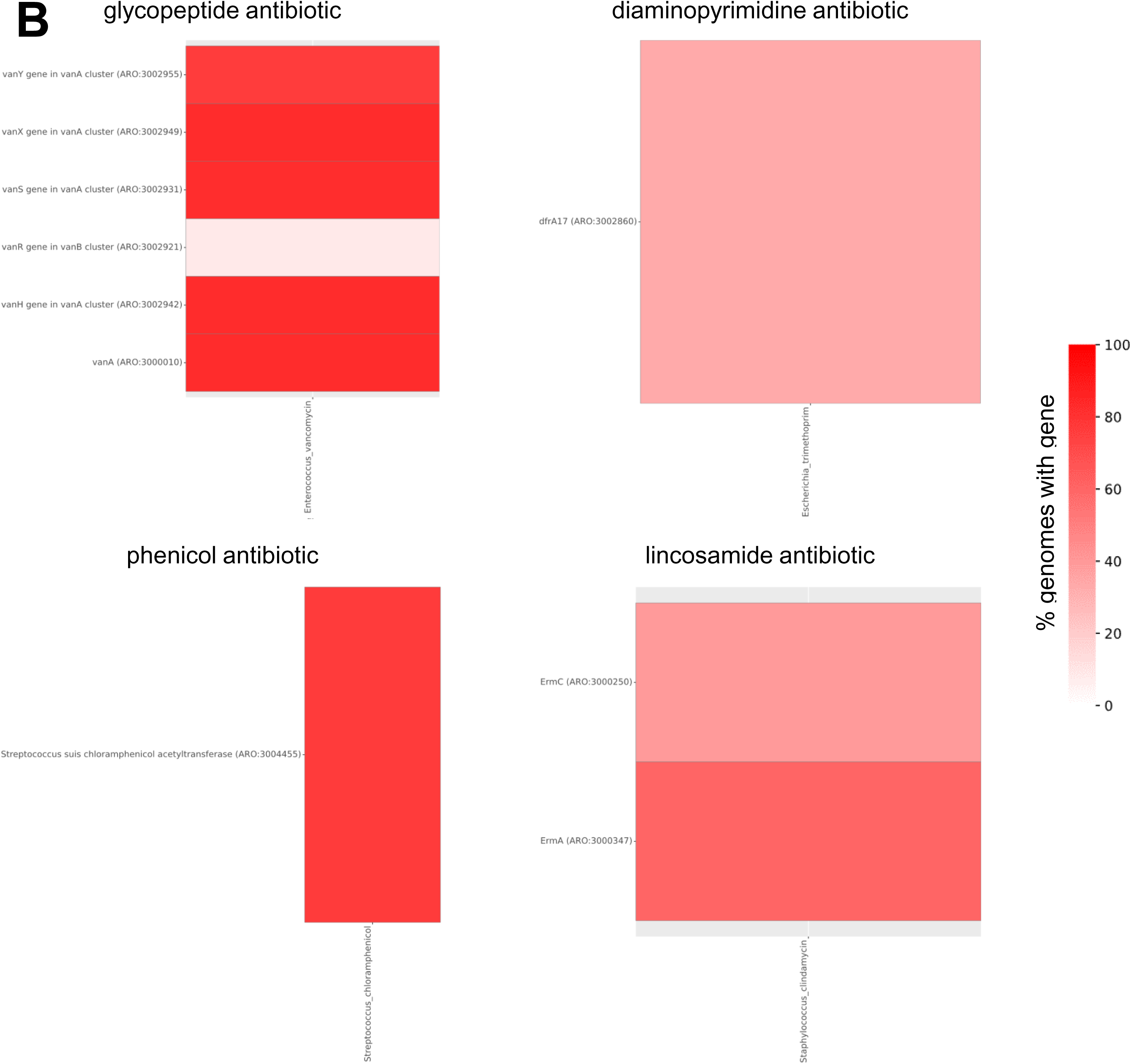
Heatmaps of known AMR gene abundance. Only genes present in ≥5% of genomes are shown.

**Figure 5.**
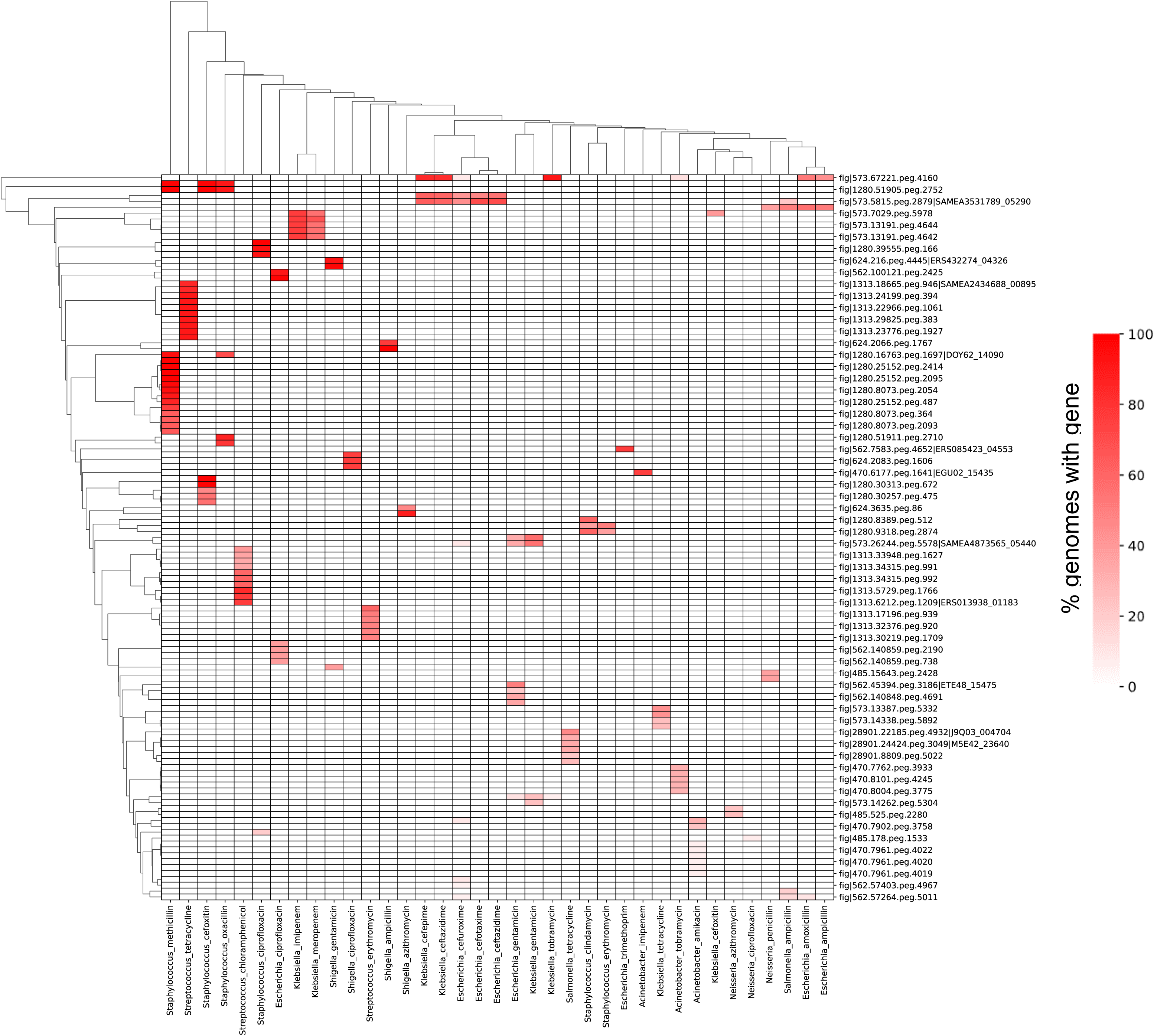
Clustermap (by Euclidean distance) of novel AMR-associated gene abundance. Only genes present in ≥5% of genomes from a pathogen-antibiotic combination are shown.

### Off-Target AMR Gene Profiles

In addition to known AMR genes conferring resistance to the target antibiotic, off-target AMR genes were identified in 23/39 pathogen-antibiotic combinations, suggesting the potential for multi-drug resistance phenotypes. Off-target AMR genes functioning as efflux pumps or modulators of drug permeability were the most often identified among different pathogen-antibiotic combinations (**Figure 6**). Chi-square tests of the co-association between on- and off-target AMR gene identified 32 gene pairs significantly associated when analyzing pairs found in >5% of genome (p-value <0.01, odds-ratio >1), in which efflux pumps were the most commonly co-harbored mechanism of action found in 8/32 gene pairs (**Table 3**).

**Figure 6:**
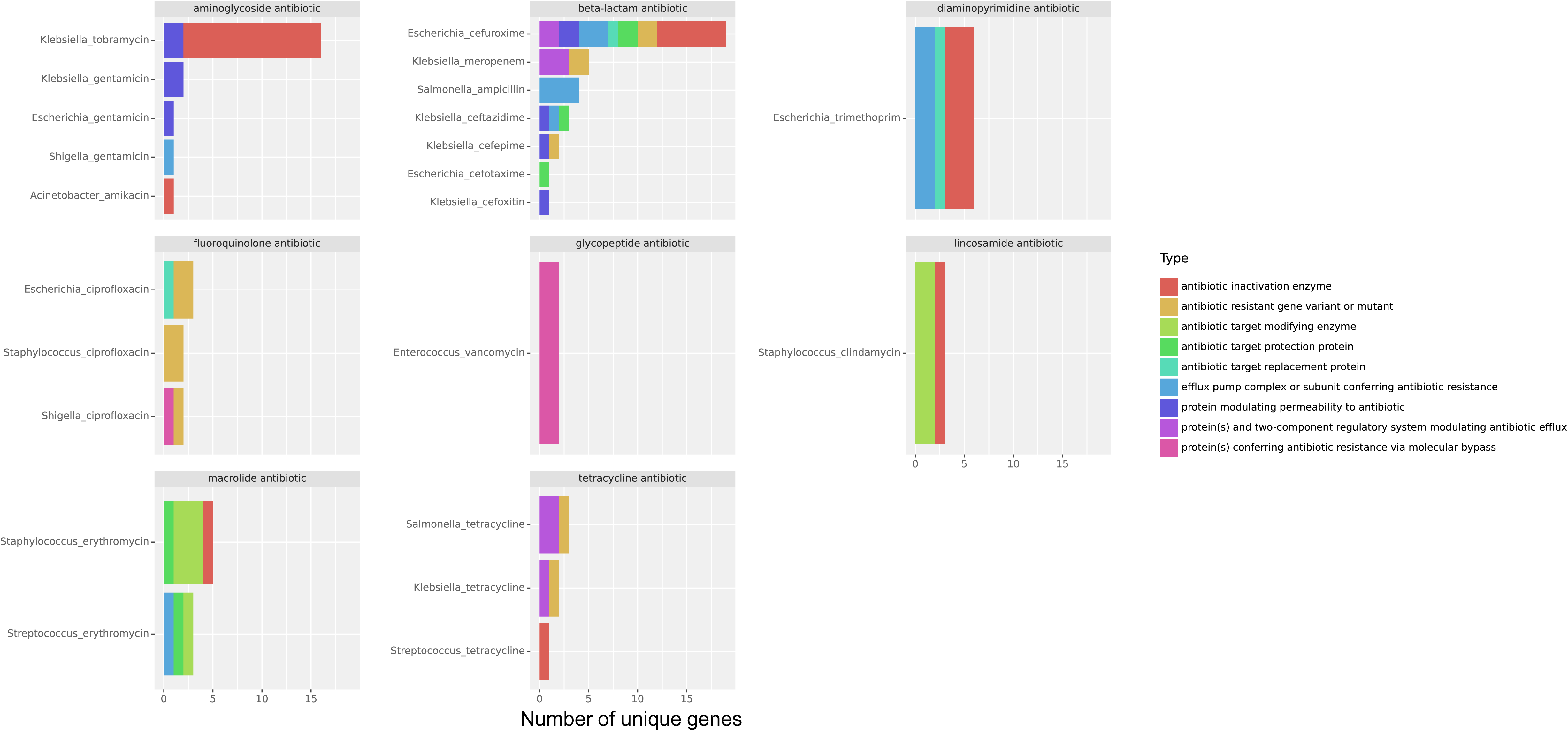
Number of unique off-target AMR genes and mechanism of action.

**Table 3.** Characteristics of Statistically Co-Association of On- with Off-Target AMR Gene Pairs.

| Pathogen-Antibiotic Combination | Gene 1 | Gene 1 MOA | Gene 1 Antibiotic Target | Gene 2 | Gene 2 MOA | Gene 2 Antibiotic Target | % genomes | Odds ratio | FDR-adjusted P-value |
| --- | --- | --- | --- | --- | --- | --- | --- | --- | --- |
| *Acinetobacter-amikacin* | armA (ARO:3000858) | rRNA methyltransferase conferring antibiotic resistance | aminoglycoside | mphE (ARO:3003741) | macrolide inactivation enzyme | macrolide antibiotic | 30.5 | inf | 3.62E-93 |
| *Enterococcus-vancomycin* | vanA (ARO:3000010) | glycopeptide resistance cluster | glycopeptide | vanR gene in vanA cluster (ARO:3002919) | glycopeptide resistance cluster | peptide antibiotic | 82.1 | 43848 | 2.37E-260 |
| *Enterococcus-vancomycin* | vanS gene in vanA cluster (ARO:3002931) | glycopeptide resistance cluster | glycopeptide | vanA (ARO:3000010) | glycopeptide resistance cluster | cycloserine-like antibiotic | 82.1 | 43848 | 2.37E-260 |
| *Enterococcus-vancomycin* | vanH gene in vanA cluster (ARO:3002942) | glycopeptide resistance cluster | glycopeptide | vanA (ARO:3000010) | glycopeptide resistance cluster | cycloserine-like antibiotic | 82.5 | inf | 2.91E-267 |
| *Enterococcus-vancomycin* | vanX gene in vanA cluster (ARO:3002949) | glycopeptide resistance cluster | glycopeptide | vanA (ARO:3000010) | glycopeptide resistance cluster | cycloserine-like antibiotic | 82.5 | inf | 2.91E-267 |
| *Enterococcus-vancomycin* | vanY gene in vanA cluster (ARO:3002955) | glycopeptide resistance cluster | glycopeptide | vanA (ARO:3000010) | glycopeptide resistance cluster | cycloserine-like antibiotic | 76.2 | 2645.299 | 7.17E-183 |
| *Enterococcus-vancomycin* | vanS gene in vanA cluster (ARO:3002931) | glycopeptide resistance cluster | glycopeptide | vanR gene in vanA cluster (ARO:3002919) | glycopeptide resistance cluster | peptide antibiotic | 82.1 | inf | 1.87E-268 |
| *Enterococcus-vancomycin* | vanH gene in vanA cluster (ARO:3002942) | glycopeptide resistance cluster | glycopeptide | vanR gene in vanA cluster (ARO:3002919) | glycopeptide resistance cluster | peptide antibiotic | 82.1 | inf | 1.01E-261 |
| *Enterococcus-vancomycin* | vanX gene in vanA cluster (ARO:3002949) | glycopeptide resistance cluster | glycopeptide | vanR gene in vanA cluster (ARO:3002919) | glycopeptide resistance cluster | peptide antibiotic | 82.1 | inf | 1.01E-261 |
| *Enterococcus-vancomycin* | vanY gene in vanA cluster (ARO:3002955) | glycopeptide resistance cluster | glycopeptide | vanR gene in vanA cluster (ARO:3002919) | glycopeptide resistance cluster | peptide | 76.1 | 911.7244 | 1.18E-182 |
| *Escherichia-cefuroxime* | CTX-M-15 (ARO:3001878) | beta-lactamase | beta-lactam | catB3 (ARO:3002676) | chloramphenicol acetyltransferase (CAT) | phenicol | 15.0 | 137.1058 | 1.24E-50 |
| *Escherichia-cefuroxime* | CTX-M-15 (ARO:3001878) | beta-lactamase | beta-lactam | catB3 (ARO:3002676) | chloramphenicol acetyltransferase (CAT) | streptogramin | 15.0 | 137.1058 | 1.24E-50 |
| *Escherichia-ciprofloxacin* | *E. coli* gyrA (ARO:3003294) | antibiotic resistant DNA topoisomerase subunit | fluoroquinolone | *E. coli* parC (ARO:3003308) | antibiotic resistant DNA topoisomerase subunit | disinfecting agents and antiseptics | 84.7 | 112.1707 | 4.01E-69 |
| *Escherichia-ciprofloxacin* | *E. coli* parC (ARO:3003308) | antibiotic resistant DNA topoisomerase subunit | fluoroquinolone | *E. coli* gyrA (ARO:3003294) | antibiotic resistant DNA topoisomerase subunit | disinfecting agents and antiseptics | 84.7 | 112.1707 | 4.01E-69 |
| *Escherichia-gentamicin* | AAC(3)-Ile (ARO:3004621) | aminoglycoside modifying enzyme | aminoglycoside antibiotic | tmrB (ARO:3003059) | tunicamycin resistance protein | nucleoside antibiotic | 34.3 | 150.8684 | 1.37E-47 |
| *Escherichia-trimethoprim* | dfrA17 (ARO:3002860) | antibiotic resistant dihydrofolate reductase | diaminopyrimidine | qacEdelta1 (ARO:3005010) | small multidrug resistance (SMR) antibiotic efflux pump | macrolide antibiotic | 30.3 | 21.10884 | 5.07E-32 |
| *Escherichia-trimethoprim* | dfrA17 (ARO:3002860) | antibiotic resistant dihydrofolate reductase | diaminopyrimidine | qacEdelta1 (ARO:3005010) | small multidrug resistance (SMR) antibiotic efflux pump | aminocoumarin antibiotic | 30.3 | 21.10884 | 5.07E-32 |
| *Escherichia-trimethoprim* | dfrA17 (ARO:3002860) | antibiotic resistant dihydrofolate reductase | diaminopyrimidine | qacEdelta1 (ARO:3005010) | small multidrug resistance (SMR) antibiotic efflux pump | aminoglycoside antibiotic | 30.3 | 21.10884 | 5.07E-32 |
| *Escherichia-trimethoprim* | dfrA17 (ARO:3002860) | antibiotic resistant dihydrofolate reductase | diaminopyrimidine | qacEdelta1 (ARO:3005010) | small multidrug resistance (SMR) antibiotic efflux pump | disinfecting agents and antiseptics | 30.3 | 21.10884 | 5.07E-32 |
| *Escherichia-trimethoprim* | dfrA17 (ARO:3002860) | antibiotic resistant dihydrofolate reductase | diaminopyrimidine | aadA5 (ARO:3002605) | aminoglycoside modifying enzyme | aminoglycoside antibiotic | 32.6 | 2317.714 | 2.08E-94 |
| *Escherichia-trimethoprim* | sul1 (ARO:3000410) | sulfonamide resistant sul | sulfonamide antibiotic | dfrA17 (ARO:3002860) | antibiotic resistant dihydrofolate reductase | diaminopyrimidine antibiotic | 30.1 | 19.56571 | 2.04E-31 |
| *Escherichia-trimethoprim* | sul1 (ARO:3000410) | sulfonamide resistant sul | sulfone antibiotic | dfrA17 (ARO:3002860) | antibiotic resistant dihydrofolate reductase | diaminopyrimidine antibiotic | 30.1 | 19.56571 | 2.04E-31 |
| *Klebsiella-gentamicin* | AAC(3)-Ile (ARO:3004621) | aminoglycoside modifying enzyme | aminoglycoside antibiotic | tmrB (ARO:3003059) | tunicamycin resistance protein | nucleoside antibiotic | 33.3 | 2.025131 | 9.74E-11 |
| *Klebsiella-gentamicin* | AAC(3)-IId (ARO:3004623) | aminoglycoside modifying enzyme | aminoglycoside antibiotic | tmrB (ARO:3003059) | tunicamycin resistance protein | nucleoside antibiotic | 18.7 | 22.88775 | 2.81E-60 |
| *Klebsiella-tetracycline* | tet(A) (ARO:3000165) | major facilitator superfamily (MFS) antibiotic efflux pump | tetracycline antibiotic | tetR (ARO:3003479) | mutant efflux regulatory protein conferring antibiotic resistance | - | 45.8 | inf | 2.46E-73 |
| *Klebsiella-tetracycline* | tet(D) (ARO:3000168) | major facilitator superfamily (MFS) antibiotic efflux pump | tetracycline antibiotic | tetR (ARO:3003479) | mutant efflux regulatory protein conferring antibiotic resistance | - | 28.1 | 30.16274 | 5.82E-31 |
| *Klebsiella-tobramycin* | aadA (ARO:3002601) | aminoglycoside modifying enzyme | aminoglycoside antibiotic | AAC(6′)-Ib10 (ARO:3002581) | aminoglycoside modifying enzyme | fluoroquinolone antibiotic | 40.1 | 4.190979 | 7.90E-21 |
| *Klebsiella-tobramycin* | AAC(6′)-Ib10 (ARO:3002581) | aminoglycoside modifying enzyme | aminoglycoside antibiotic | catB3 (ARO:3002676) | chloramphenicol acetyltransferase (CAT) | phenicol antibiotic | 38.4 | 7.101639 | 1.97E-30 |
| *Klebsiella-tobramycin* | AAC(6′)-Ib10 (ARO:3002581) | aminoglycoside modifying enzyme | aminoglycoside antibiotic | catB3 (ARO:3002676) | chloramphenicol acetyltransferase (CAT) | streptogramin antibiotic | 38.4 | 7.101639 | 1.97E-30 |
| *Klebsiella-tobramycin* | AAC(6′)-Ib10 (ARO:3002581) | aminoglycoside modifying enzyme | aminoglycoside antibiotic | tmrB (ARO:3003059) | tunicamycin resistance protein | nucleoside antibiotic | 29.4 | 3.80863 | 3.79E-14 |
| *Klebsiella-tobramycin* | AAC(3)-Ile (ARO:3004621) | aminoglycoside modifying enzyme | aminoglycoside antibiotic | AAC(6′)-Ib10 (ARO:3002581) | aminoglycoside modifying enzyme | fluoroquinolone antibiotic | 28.3 | 2.626489 | 1.86E-08 |
| *Klebsiella-tobramycin* | AAC(6′)-Ib10 (ARO:3002581) | aminoglycoside modifying enzyme | aminoglycoside antibiotic | OXA-1 (ARO:3001396) | beta-lactamase | beta-lactam antibiotic | 38.9 | 8.496912 | 5.86E-34 |
| *Klebsiella-tobramycin* | AAC(3)-Ile (ARO:3004621) | aminoglycoside modifying enzyme | aminoglycoside antibiotic | OXA-1 (ARO:3001396) | beta-lactamase | beta-lactam antibiotic | 25.9 | 11.4832 | 2.07E-69 |
| *Klebsiella-tobramycin* | AAC(3)-Ile (ARO:3004621) | aminoglycoside modifying enzyme | aminoglycoside antibiotic | catB3 (ARO:3002676) | chloramphenicol acetyltransferase (CAT) | phenicol antibiotic | 25.7 | 10.83711 | 5.99E-67 |
| *Klebsiella-tobramycin* | AAC(3)-Ile (ARO:3004621) | aminoglycoside modifying enzyme | aminoglycoside antibiotic | catB3 (ARO:3002676) | chloramphenicol acetyltransferase (CAT) | streptogramin antibiotic | 25.7 | 10.83711 | 5.99E-67 |
| *Klebsiella-tobramycin* | AAC(3)-Ile (ARO:3004621) | aminoglycoside modifying enzyme | aminoglycoside antibiotic | tmrB (ARO:3003059) | tunicamycin resistance protein | nucleoside antibiotic | 27.8 | 57.90865 | 1.48E-146 |
| *Klebsiella-tobramycin* | AAC(3)-IId (ARO:3004623) | aminoglycoside modifying enzyme | aminoglycoside antibiotic | tmrB (ARO:3003059) | tunicamycin resistance protein | nucleoside antibiotic | 6.1 | 59.77536 | 2.97E-31 |
| *Salmonella-ampicillin* | CMY-2 (ARO:3002013) | beta-lactamase | beta-lactam antibiotic | ykkC (ARO:3003063) | subunit of efflux pump conferring antibiotic resistance | aminoglycoside antibiotic | 17.3 | 2816.929 | 5.31E-240 |
| *Salmonella-ampicillin* | CMY-2 (ARO:3002013) | beta-lactamase | beta-lactam antibiotic | ykkC (ARO:3003063) | small multidrug resistance (SMR) antibiotic efflux pump | aminoglycoside antibiotic | 17.3 | 2816.929 | 5.31E-240 |
| *Salmonella-tetracycline* | tet(A) (ARO:3000165) | major facilitator superfamily (MFS) antibiotic efflux pump | tetracycline antibiotic | tetR (ARO:3003479) | mutant efflux regulatory protein conferring antibiotic resistance | - | 55.6 | 136.0778 | 6.41E-49 |
| *Salmonella-tetracycline* | tet(B) (ARO:3000166) | major facilitator superfamily (MFS) antibiotic efflux pump | tetracycline antibiotic | tetR (ARO:3003479) | mutant efflux regulatory protein conferring antibiotic resistance | - | 33.6 | 10.41443 | 7.86E-16 |
| *Salmonella-tetracycline* | tet(B) (ARO:3000166) | major facilitator superfamily (MFS) antibiotic efflux pump | tetracycline antibiotic | adeN (ARO:3000559) | resistance-nodulation-cell division antibiotic efflux pump | - | 12.0 | 239.2074 | 6.72E-76 |
| *Staphylococcus-ciprofloxacin* | *S. aureus* gyrA (ARO:3003296) | antibiotic resistant DNA topoisomerase subunit | fluoroquinolone antibiotic | *S. aureus* parC (ARO:3003312) | antibiotic resistant DNA topoisomerase subunit | disinfecting agents and antiseptics | 90.5 | 18.55357 | 1.49E-35 |
| *Staphylococcus-ciprofloxacin* | *S. aureus* parC (ARO:3003312) | antibiotic resistant DNA topoisomerase subunit | fluoroquinolone antibiotic | *S. aureus* gyrA (ARO:3003296) | antibiotic resistant DNA topoisomerase subunit | disinfecting agents and antiseptics | 90.5 | 18.55357 | 1.49E-35 |
| *Staphylococcus-clindamycin* | ErmA (ARO:3000347) | rRNA methyltransferase conferring antibiotic resistance | lincosamide antibiotic | ANT(9)-Ia (ARO:3002630) | aminoglycoside modifying enzyme | aminoglycoside antibiotic | 60.2 | inf | 1.16E-100 |
| *Staphylococcus-erythromycin* | ErmA (ARO:3000347) | rRNA methyltransferase conferring antibiotic resistance | macrolide antibiotic | ANT(9)-Ia (ARO:3002630) | aminoglycoside modifying enzyme | aminoglycoside antibiotic | 37.5 | inf | 5.55E-198 |
| *Streptococcus-erythromycin* | mel (ARO:3000616) | major facilitator superfamily (MFS) antibiotic efflux pump | macrolide antibiotic | msrE (ARO:3003109) | ABC-F ATP-binding cassette ribosomal protection protein | phosphonic acid antibiotic | 51.9 | inf | 0 |
| *Streptococcus-erythromycin* | mel (ARO:3000616) | major facilitator superfamily (MFS) antibiotic efflux pump | macrolide antibiotic | msrE (ARO:3003109) | ABC-F ATP-binding cassette ribosomal protection protein | streptogramin antibiotic | 51.9 | inf | 0 |
| *Streptococcus-erythromycin* | mel (ARO:3000616) | major facilitator superfamily (MFS) antibiotic efflux pump | macrolide antibiotic | msrE (ARO:3003109) | ABC-F ATP-binding cassette ribosomal protection protein | lincosamide antibiotic | 51.9 | inf | 0 |
| *Streptococcus-erythromycin* | mel (ARO:3000616) | major facilitator superfamily (MFS) antibiotic efflux pump | macrolide antibiotic | msrE (ARO:3003109) | ABC-F ATP-binding cassette ribosomal protection protein | pleuromutilin antibiotic | 51.9 | inf | 0 |
| *Streptococcus-erythromycin* | mel (ARO:3000616) | major facilitator superfamily (MFS) antibiotic efflux pump | macrolide antibiotic | msrE (ARO:3003109) | ABC-F ATP-binding cassette ribosomal protection protein | nucleoside antibiotic | 51.9 | inf | 0 |
| *Streptococcus-erythromycin* | mel (ARO:3000616) | major facilitator superfamily (MFS) antibiotic efflux pump | macrolide antibiotic | msrE (ARO:3003109) | ABC-F ATP-binding cassette ribosomal protection protein | antibiotic without defined classification | 51.9 | inf | 0 |
| *Streptococcus-erythromycin* | msrE (ARO:3003109) | ABC-F ATP-binding cassette ribosomal protection protein | macrolide antibiotic | mel (ARO:3000616) | major facilitator superfamily (MFS) antibiotic efflux pump | tetracycline antibiotic | 51.9 | inf | 0 |

### Functional Domain Predictions of Novel AMR-Associated Genes

To gain the functional insights of “novel” AMR-associated genes, protein domains were predicted using InterProScan. Protein domains were identified in 4,001/5,224 (70.8%) of the novel AMR-associated gene sequence. A total of 901 unique domains were present, representing a wide diversity of potential functions and mechanisms. Hierarchal cluster analysis indicated the types and distribution of proteins domains tended to associate with more similar pathogen-antibiotic combinations, which was also observed for known on- and off-target AMRs genes as described above (**Figure 7**). Across pathogen-antibiotic combinations, the most commonly observed domains were related to horizonal gene transfer and included plasmid- and phage-mediated genetic exchange mechanisms such as ribonuclease, resolvase, integrase, recombinase, transposase, endonuclease proteins. This finding supports the importance of genetic exchange being a critical mechanism driving the propagation of antibiotic resistance in pathogens. Aside from horizonal gene transfer, domains implicated in molecular transporters and cell wall structure were frequently observed, which may expand the repertoire of efflux pumps/modulators of membrane permeability that contribute to resistance (**Figure 8, 9**). Interestingly, several domains related to oxidation-reduction mechanisms, such as FAD-linked oxidoreductase, NADP-dependent oxidoreductase, and flavoprotein-like domains were identified (**Figure 8, 9**). Domains associated with antibiotic/natural product metabolism including nonribosomal peptide synthetase, antibiotic biosynthesis monooxygenase, isochorismatase-like hydrolase, chloramphenicol acetyltransferase-like, dihydroxybiphenyl dioxygenase were also identified (**Figure 8, 9**). These protein domains may contribute to antibiotic resistance via degradation mechanisms. Notably, several domains similar to known AMR mechanism like dihydrofolate reductase subunits, pentapeptide repeat-like, DNA topoisomerase, tetracycline repressor-like and were detected as well (**Figure 8, 9**).

**Figure 7.**
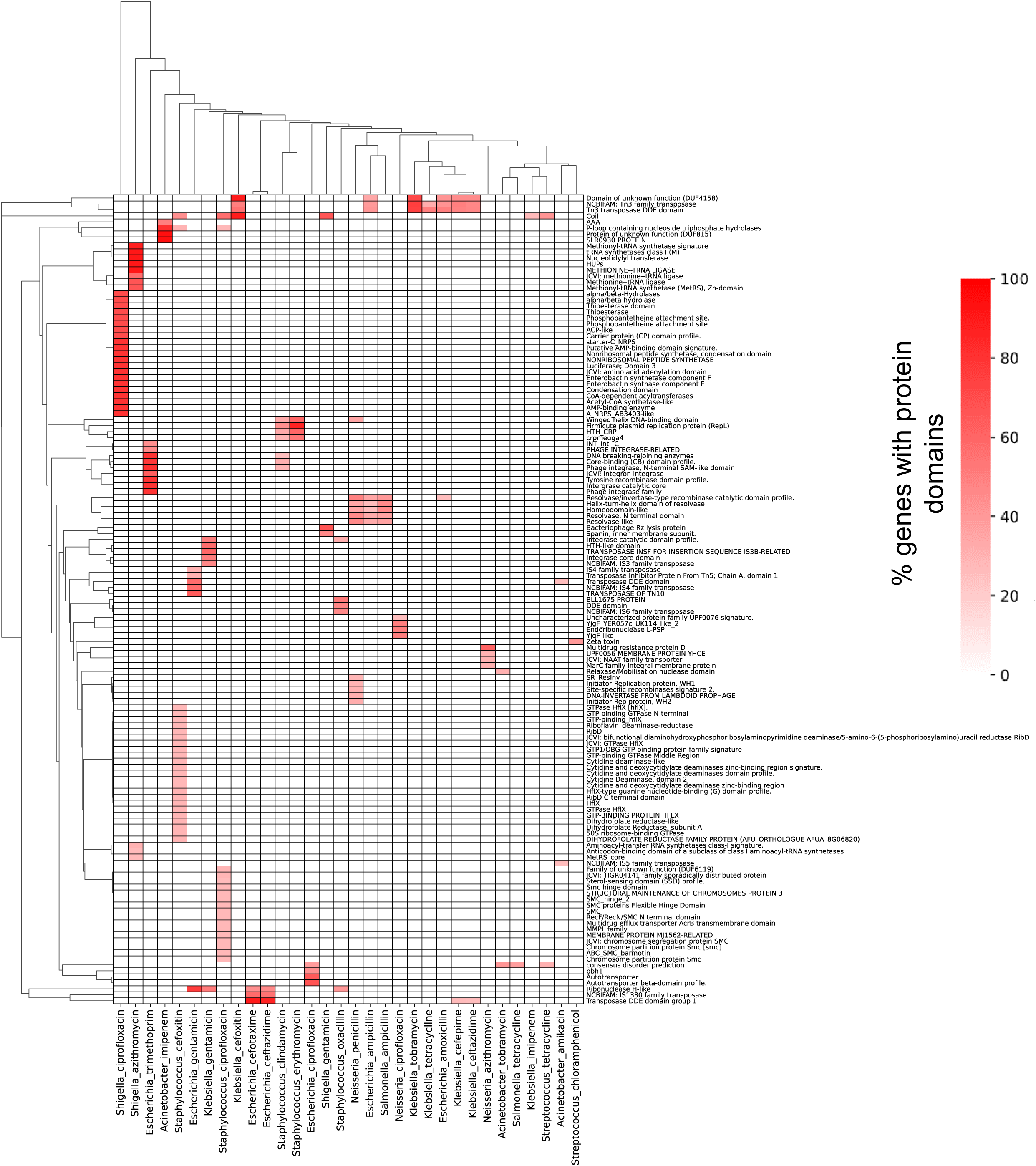
Clustermap (by Euclidean distance) of protein domains abundance in novel AMR - associated genes. Only protein domains present in ≥25% of genes for a pathogen-antibiotic combination are shown.

**Figure 8.**
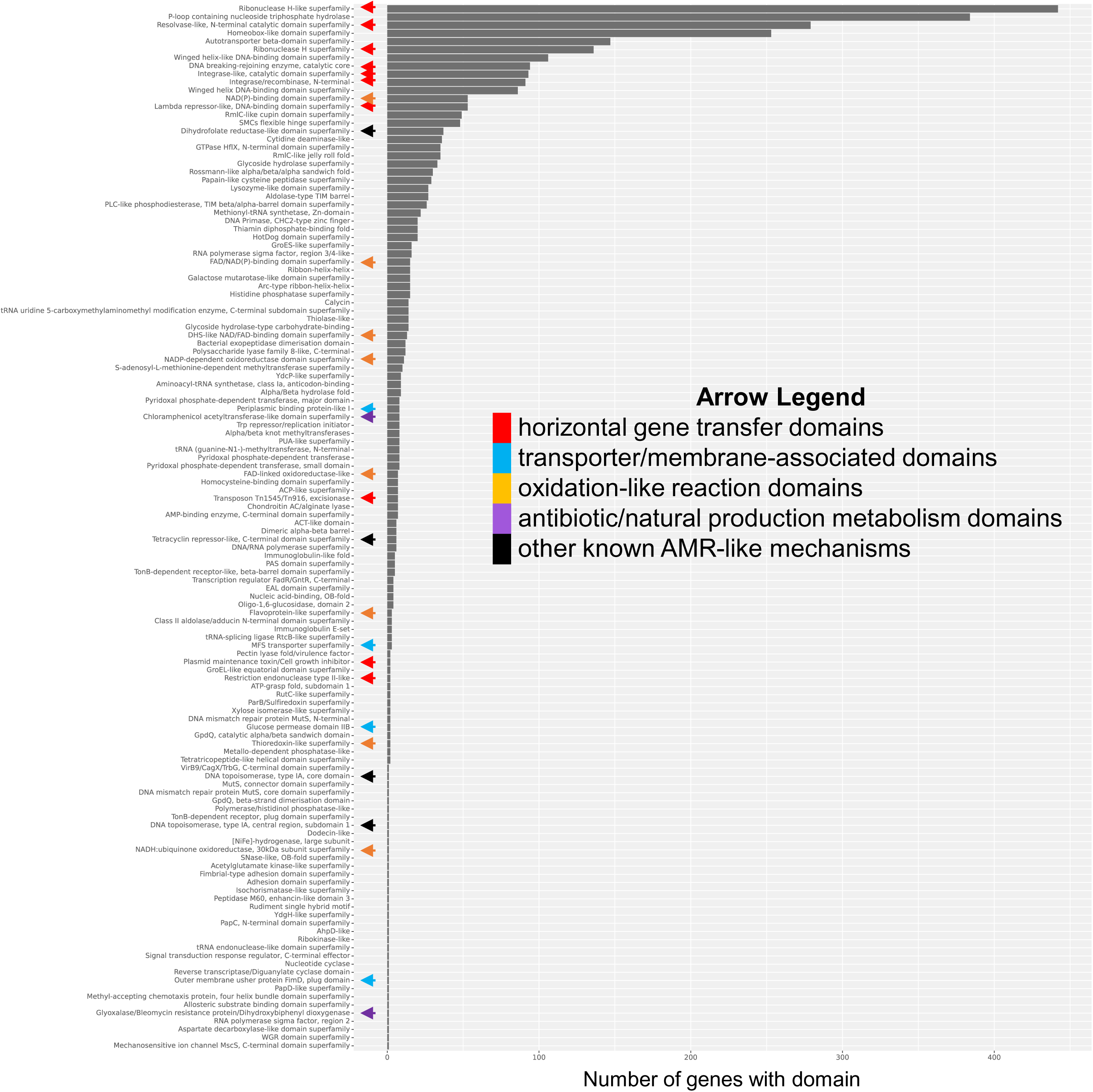
Absolute abundance of protein domain superfamilies (at InterPro description level) in novel associated genes.

**Figure 9.**
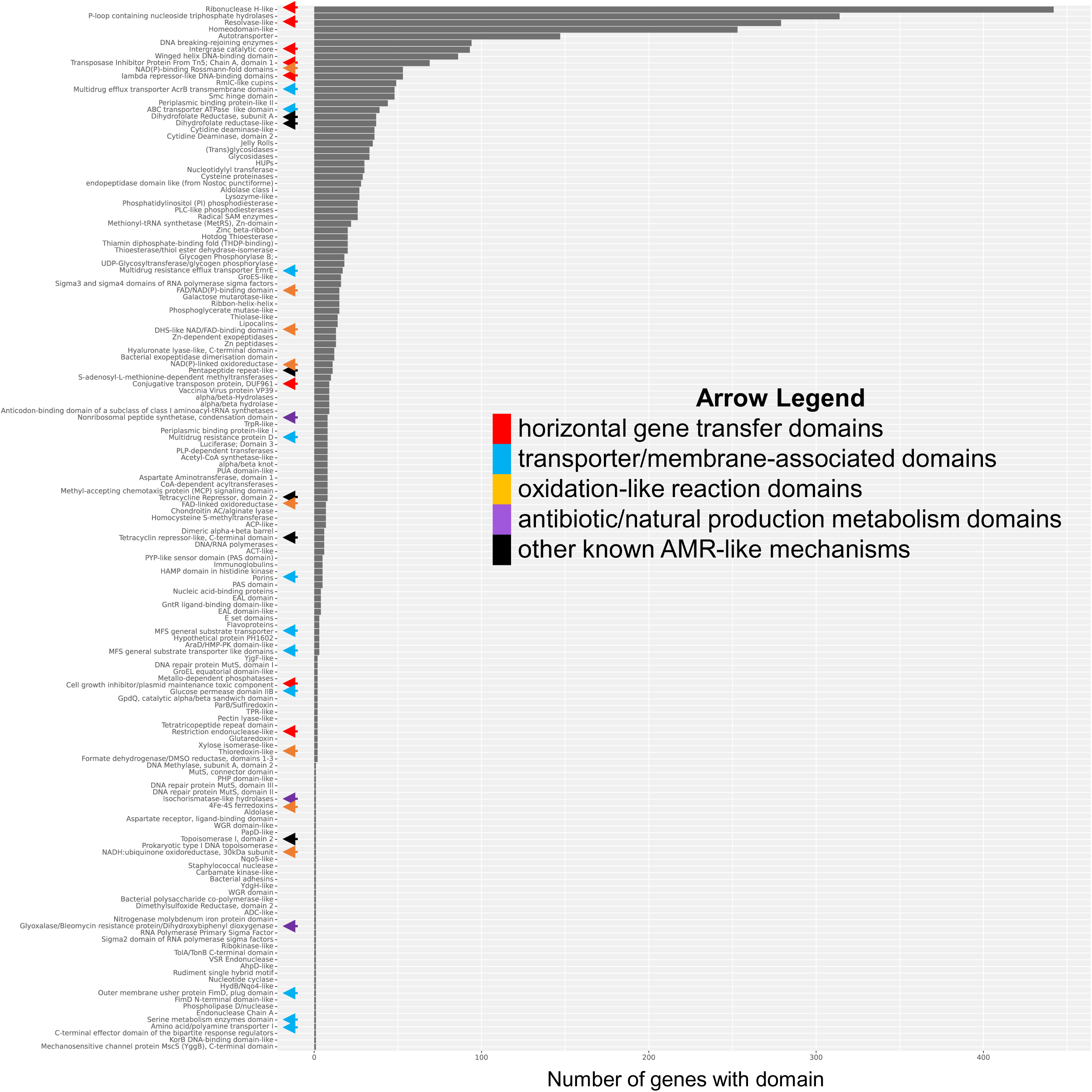
Absolute abundance of protein domain superfamilies (at signature description level) in novel associated genes.

### Structure-Function Modeling of Novel AMR-Associated Genes

Random Forest and XGboost models were created to classify protein-antibiotic interactions based on predicted protein-ligand binding affinity profiles, protein sequence embeddings and antibiotic molecular fingerprints/descriptors. As shown in the parallel coordinate plots for standardized and non-standardized performance metrics in (**Figure 10**), the Random Forest model achieved higher precision, specificity, AUC, and accuracy values compared to XGboost. PRAUC scores were comparable between the models, while XGboost achieved higher sensitivity (**Figure 10**). Both models yielded better performance on macrolides compared to other antibiotic classes (**Figure 10**). Aminoglycosides and beta-lactams had higher number of estimated interactions with lower specificity and accuracy but higher sensitivity.

**Figure 10.**
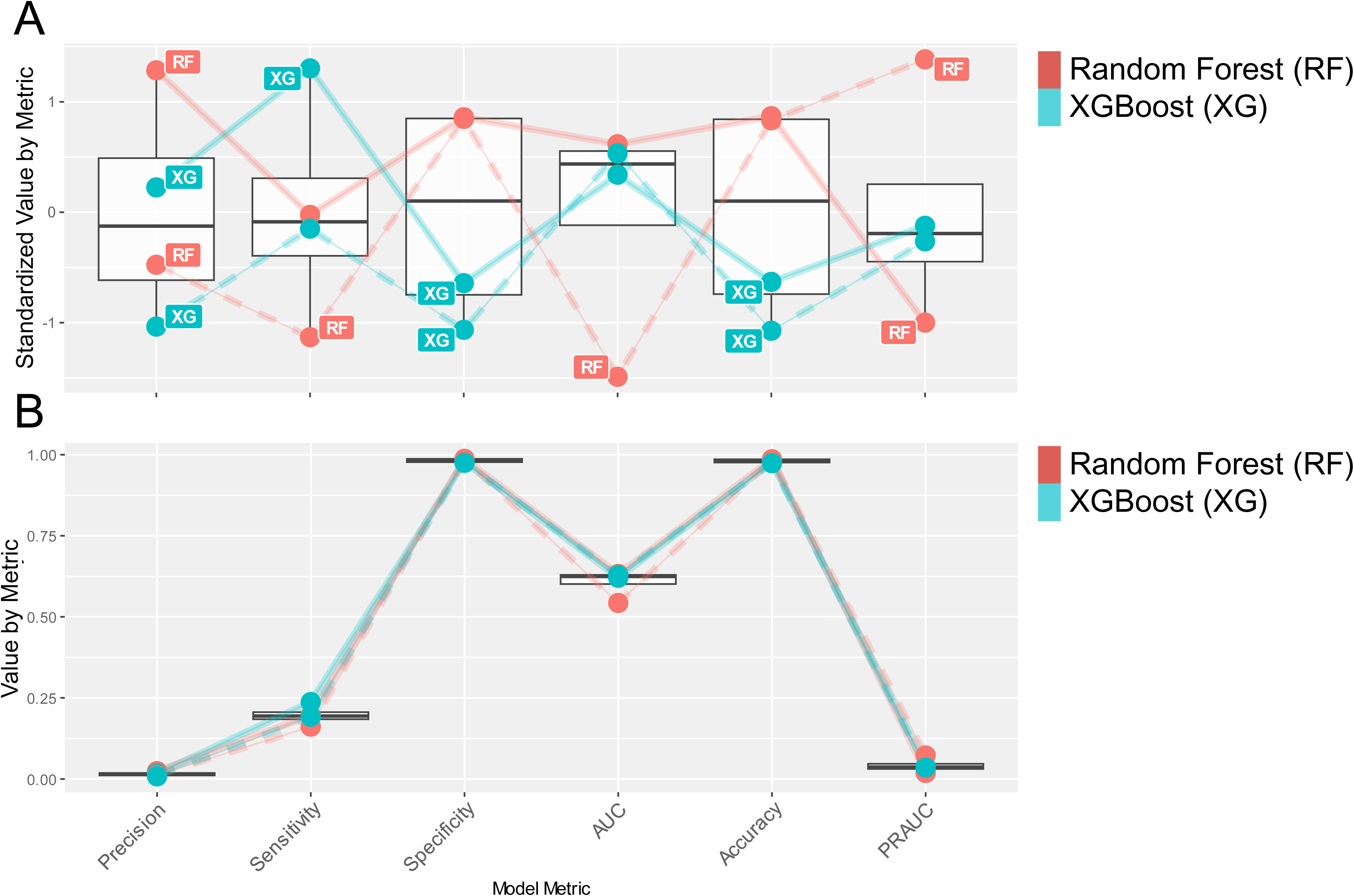
Comparison of standardized (A) and non-standardized (B) model metrics Random Forest (RF) and XGBoost (XG) classifiers. Solid lines represent model performance over all values and dashed lines represent average model performance across antibiotic class.

Additionally, both models were assessed on different data subsets based on whether the proteins and ligands were present or not in the training dataset to provide more insights into the model performance. The subsets evaluated were: S1) both proteins and ligands not in training (**Supplemental Figure 3**), S2) proteins in training and ligands not in training (**Supplemental Figure 4**), and S3) ligands in training and proteins not in training (**Supplemental Figure 5)**. Analysis on S1 and S3 show that the model is able to identify protein-ligand interactions for specific antibiotics not used in training but belonging to an antibiotic class used in training. This suggests protein interactions with new molecular derivatives of existing antibiotic classes could be identified. Importantly, both models exhibited higher precision, specificity, AUC, and PRAUC scores for S3 versus S1 or S2, suggesting this approach can identify potential protein-ligand interaction for novel, unseen proteins (**Figure 11**). Based on overall performance characteristics between the two models, the Random Forest classifier was chosen to characterize the novel AMR-associated protein for interactions with antibiotics.

**Figure 11.**
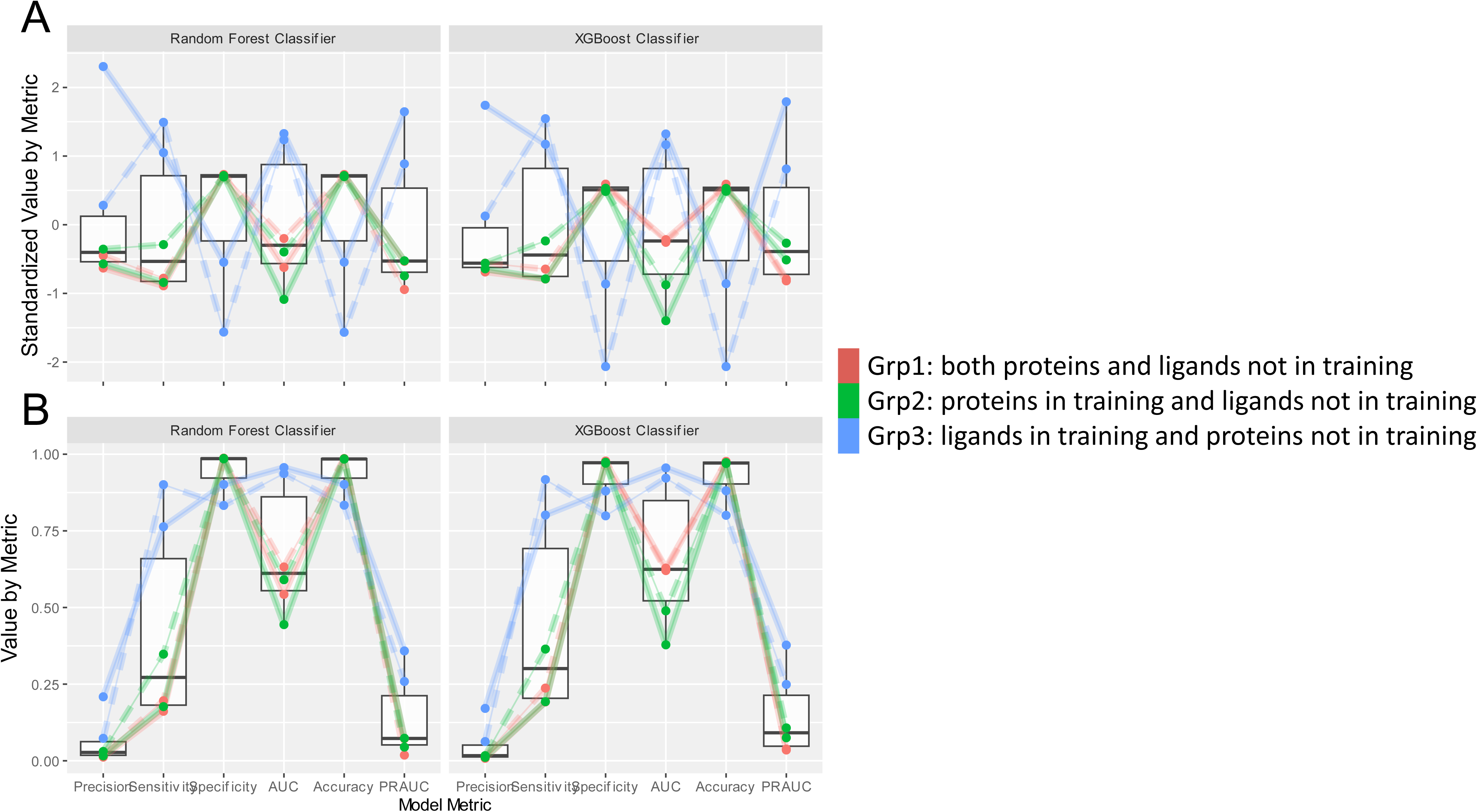
Comparison of standardized (A) and non-standardized (B) model metrics Random Forest (RF) and XGBoost (XG) classifiers on Grp1 (both proteins and ligands not in training), Grp2 (proteins in training and ligands not in training), and Grp3 (ligands in training and proteins not in training) data subsets. Solid lines represent model performance over all values and dashed lines represent average model performance across antibiotic class.

Using the random forest model, 45 protein-antibiotic combinations were identified with interaction probabilities >0.5. Of these, 15 proteins with amino acid length >100 residues were considered for further analysis (**Table 4**). Only protein-antibiotic interactions for aminoglycosides were identified in the hits. However, several protein sequences were also found in *Klebsiella* strains resistant to cefepime and ceftazidime (**Figure 12**). Interestingly, protein hits had a single or no protein domains annotated based on InterProScan analysis. The most common domain identified was “Domain of unknown function (DUF4158)” found in 8/15 of the hits. 3/15 proteins had no protein domains identified, suggesting uncharacterized structure and function. Of the 15 proteins identified, 9 belonged to the same cluster (based on >30% sequence identify).

**Figure 12.**
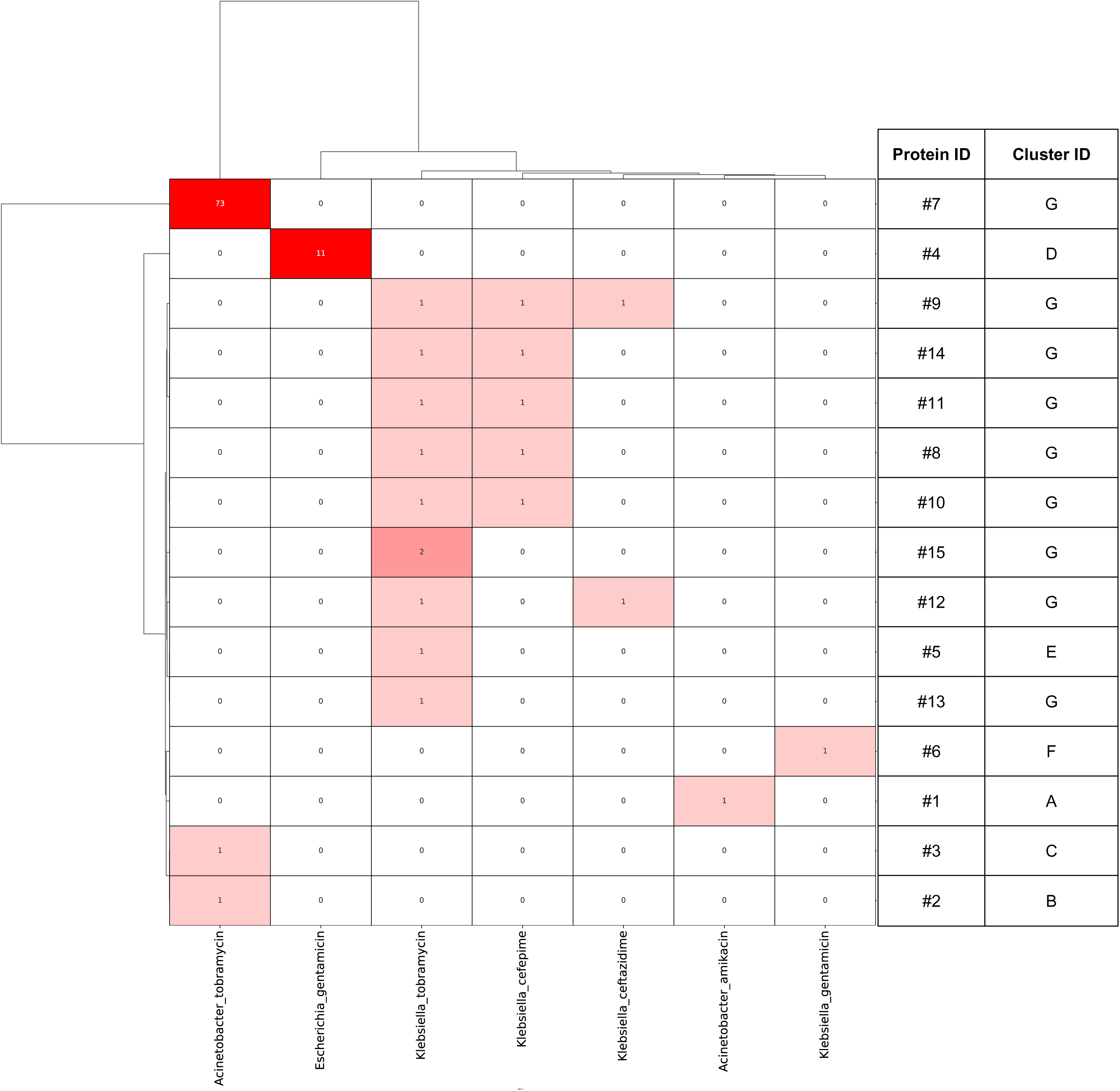
Clustermap (by Euclidean distance) showing number of genomes per pathogen-antibiotic combinations containing protein with identified ligand interactions. Number of genomes containing gene are indicated within cells.

**Table 4:**
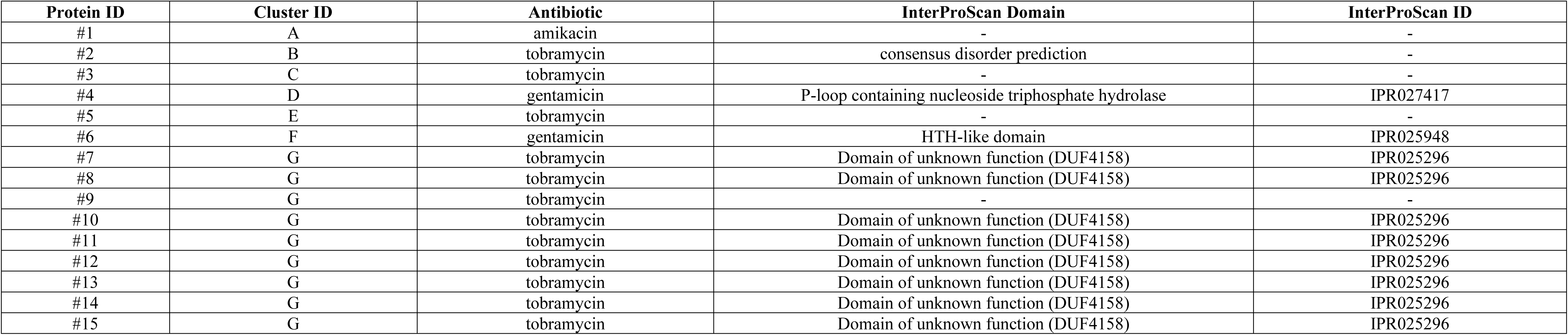
Proteins identified with ligand interactions.

| Protein ID | Cluster ID | Antibiotic | InterProScan Domain | InterProScan ID |
| --- | --- | --- | --- | --- |
| #1 | A | amikacin | - | - |
| #2 | B | tobramycin | consensus disorder prediction | - |
| #3 | C | tobramycin | - | - |
| #4 | D | gentamicin | P-loop containing nucleoside triphosphate hydrolase | IPR027417 |
| #5 | E | tobramycin | - | - |
| #6 | F | gentamicin | HTH-like domain | IPR025948 |
| #7 | G | tobramycin | Domain of unknown function (DUF4158) | IPR025296 |
| #8 | G | tobramycin | Domain of unknown function (DUF4158) | IPR025296 |
| #9 | G | tobramycin | - | - |
| #10 | G | tobramycin | Domain of unknown function (DUF4158) | IPR025296 |
| #11 | G | tobramycin | Domain of unknown function (DUF4158) | IPR025296 |
| #12 | G | tobramycin | Domain of unknown function (DUF4158) | IPR025296 |
| #13 | G | tobramycin | Domain of unknown function (DUF4158) | IPR025296 |
| #14 | G | tobramycin | Domain of unknown function (DUF4158) | IPR025296 |
| #15 | G | tobramycin | Domain of unknown function (DUF4158) | IPR025296 |

## Discussion

AMR pathogens remain a significant health crisis and require improved multi-modal approaches to identify and counteract these threats. Accordingly*, The National Action Plan for Combating Antibiotic-Resistant Bacteria* (CARB) issued by the U.S. Government emphasizes the necessity to mitigate the emergence and spread of AMR pathogens by advancing therapeutic and diagnostic technologies, including approaches for biosurveillance. Traditional biosurveillance for AMR pathogens have relied on laboratory-based methods to isolate infectious agents followed by antibiotic susceptibility testing or PCR-based assays for AMR genes against limited target panels. In the last decade, the increasing adoption of whole genome sequencing has enabled holistic characterization of AMR-associated determinants in pathogens. Thus, in the current study, we performed a large-scale comparative genomic analysis of AMR pathogens to identify key trends in resistance gene profiles and augment the biosurveillance landscape.

We utilized the BV-BRC’s repository of whole genome sequences paired with laboratory-based antibiogram data to investigate over 70k ESKAPE+ pathogen strains. Our dataset included commonly used classes of antibiotics, thereby complementing the wide genomic diversity of the pathogen strains. By comparing protein-coding genes prevalence between susceptible versus resistant genome strains using a kmer-based GWAS approach, over 7,000 unique sequences were significantly enriched in resistant strains. When compared to CARD, 1,925 (29.9%) were homologous to known AMR genes, while over 5,000 (>70%) of enriched genes lacked sequence similarity. In addition to suggesting the presence of novel AMR-associated genes, this finding indicates that curated databases for AMR genes,^16^ such as CARD, may have gaps in their catalogues that could detrimentally impact the characterization of known or emerging threats.

Overall, the AMR gene profile for both known and novel-associated sequences was influenced according to pathogen-antibiotic combination. Pathogens with closer phylogeny and antibiotics with more similar molecular structures tended to have more comparable gene profiles. Additionally, we observed that only select known AMR genes predominated per pathogen-antibiotic combinations. Previous studies have described how pathogen resistomes converge on particular resistance genes/mechanisms in order to optimize survival against specific generations of beta-lactam or tetracycline antibiotics.^17^ When taken together, this suggests evolutionary pressures or niche-specific mechanisms may influence which AMR genes will prevail in pathogens. In support of this observation, horizonal gene transfer mechanisms were frequently identified as enriched AMR-associated genes, which may imply that selective exchange of AMR genes occur among more similar pathogen-antibiotic combination types. While horizonal gene transfer is well-described mechanism for AMR gene propagation, a recent study analyzing metagenomic sequencing data found that environmental niches may influence specific horizontal gene transfer mechanisms for AMR genes.^18^ Due to metadata limitation in our dataset, we were unable to investigate how environmental factors associated with AMR gene profiles. However, we hypothesize that in addition to “classic” AMR genes, tracking and characterization of horizontal gene transfer mechanisms may improve the impact of biosurveillance efforts.

Previous reports have indicated that AMR can be driven by the co-production of multiple resistance genes simultaneously. For example, *Klebsiella* and other Gram-negative enteric pathogen have been identified that harbor two or beta-lactamase genes, especially carbapenemases which target imipenem, meropenem and other carbapenems.^19^ Similarly, *Klebsiella* isolates co-harboring multiple aminoglycoside inactivation enzymes have been described.^20^ In our analysis, multi-gene resistance strains were identified across numerous antibiotic classes. However, based on statistical analysis by the chi-square tests, the co-presence of multiple for on-target antibiotic was infrequent. Aside from the vancomycin gene resistance cluster, enzymatic inactivation enzymes targeting aminoglycosides were found to be statistically co-harbored in the *Klebsiella*-tobramycin and *Salmonella*-streptomycin combinations. It is possible multi-gene resistance appear more frequently, but underrepresentation of these strains prevented statistical detection. Larger and more diverse datasets could enhance the profiling and identification of multi-gene resistance patterns in pathogens.

From our analyses, we also gained insights into the gene profiles for multidrug resistant mechanisms. We identified that 23/39 pathogen-antibiotic combination analyzed had strains with off-target AMR genes present, reinforcing the risk for multidrug resistance in pathogens. However, we only identified 32 AMR gene pairs that were significantly associated with multidrug resistant potential, with the most often occurring mechanisms being due to efflux pumps found in a diversity of pathogens. Interestingly, *Klebsiella* strains resistant to tobramycin often co-harbored inactivation enzymes for over 4 non-aminoglycoside antibiotic classes. In contrast, *Klebsiella* strains resistant to gentamicin (also an aminoglycoside) or *Escherichia* strains resistant to tobramycin did not share a diverse multidrug resistance profile. This observation suggests that different selection factors may contribute to the acquisition of multidrug resistant determinants and that these mechanisms could be specific for pathogen-antibiotic combinations.

Through our comparative genomics approach, we also identified over 5,000 unique protein sequences without homology to known AMR gene in the CARD. To investigate the potential function of these sequences, we performed protein domain analysis. As mentioned above, domains for horizonal gene transfer, such as plasmid and phage proteins, were frequently identified in the novel gene set. However, notable domains associated with antibiotic / natural product metabolism were also detected. Because most antibiotics are derived from biosynthetic gene clusters expressed by bacterial and fungal species, there are numerous examples of how these genes also contribute to resistance. For example, the “producer hypothesis” proposes that a possible origin of antibiotic resistance is due to self-protecting mechanisms towards the cytotoxic molecule being produced by source organisms.^21,22^ These enzymes may have promiscuous substrate specificity or evolve new functions that confer resistance to other molecules, and this process may occur in tandem with genetic transmission to new hosts. In the biosynthesis of natural products, microbes frequently employ enzyme-mediate oxidation reactions.^22–24^ By co-opting similar oxidation mechanisms, microbes have acquired antibiotic resistance, such as FAD-dependent oxidoreductase enzymes inactivating tetracyclines.^25,26^ In our novel gene set, several protein domains associated with oxidation activity were identified, suggesting the possibility these proteins may potentiate resistance under similar mechanisms. Furthermore, several notable protein domains associated with antibiotic-associated metabolism/resistance were detected such as isochorismatase-like hydrolase, chloramphenicol acetyltransferase-like, glyoxalase / dihydroxybiphenyl dioxygenase, which may target to streptothricin, phenicol, and bleomycin- / fosfomycin-like molecules, respectively.^27^ Likewise, known resistance gene may further mutate to accommodate non-canonical molecular scaffolds, as demonstrated by aminoglycoside N-acetyltransferase variants that can target and inactivate some fluoroquinolones.^28–30^

Our analysis also detected several proteins with domains associated with known AMR mechanisms. These included multi-drug transporter, dihydrofolate reductase-like, topoisomerase, and pentapeptide-like repeat domains. While the identification of these proteins suggests that reference AMR gene database, like CARD, may have incomplete variants catalogs, additional verification is needed to confirm the role of these proteins with resistance phenotypes. Interestingly, 1,523 (29.1%) of novel genes associated with AMR identified in our study lacked protein domains, which suggests the potential for novel mechanisms that may influence the genetic transmission of resistance mechanisms and/or potentiate antibiotic susceptibility.

Antibiotic inactivation enzymes are common mechanisms utilized to confer resistance and rely on direct interactions between protein and ligands. Thus, recent advances in AI/ML-based modeling of 3D protein structure with ligand docking could be an additional tool to identify and character AMR-associated proteins at high-throughput scales. Several previous studies have utilized structure-function modeling methods to investigate protein interactions with antibiotic targets, for both identifying potentially new antibiotic targets as well as identifying/classifying potentially new AMR-associated proteins.^31–33^ In our study, we utilized ESM-Fold and DiffDock2 to model protein structure and docking of antibiotic molecules, along with GNINA to refine docking poses and calculate binding affinity properties. To focus on the identification of antibiotic inactivation mechanisms, we trained a random forest model that also incorporated protein structure and molecular features along with binding affinity profiles based on known protein-antibiotic interactions for inactivation enzymes from the CARD. Using our random forest model, we identified several novel protein sequences that may engage with their target antibiotic molecules, including several proteins that lacked known domain regions. While further experimental work is needed to elucidate the contributions of these proteins to AMR phenotypes, our analysis approach provides a means to identify and characterize novel AMR-associated genetic determinants with statistical confidence. Furthermore, continued improvements in structure-function modeling tools along with expansions of curated AMR gene databases could enhance model performance and capabilities, such as the application of multi-classification algorithms to identify different resistance mechanism simultaneously

Despite our comprehensive comparative analysis approach, several pathogen-antibiotic combinations included genomes that lacked AMR-associated genes. It is possible our analysis approach was underpowered to detect statistically associated genes contributed by these strains. Additionally, alternative mechanisms, such as epigenetic/transcriptional control of gene expression or post-translational modifications, may promote resistance instead of acquired AMR genes.^34^ Nonetheless, our data analysis approach provides a means to identify such pathogens so that their cryptic mechanism of resistance can be further investigated.

## Conclusions

In conclusion, our findings support our hypothesis that large-scale analysis of whole genome sequence data from pathogens followed by downstream structure-function predictions are feasible approaches to identify and characterize the genetic determinants of AMR in pathogens. Continued application of and investment in these computational tools may enhance biosurveillance efforts against AMR as well as promote new means to counteract emerging threats.

## Acknowledgements

This work was funded by the Biomedical Sciences and Technologies Technical Investment Portfolio at MIT Lincoln Laboratory. The authors acknowledge the MIT Lincoln Laboratory Supercomputing Center for providing high-performance computing resources that have contributed to the research results reported herein.

---

**Supplemental Figure 1A.**
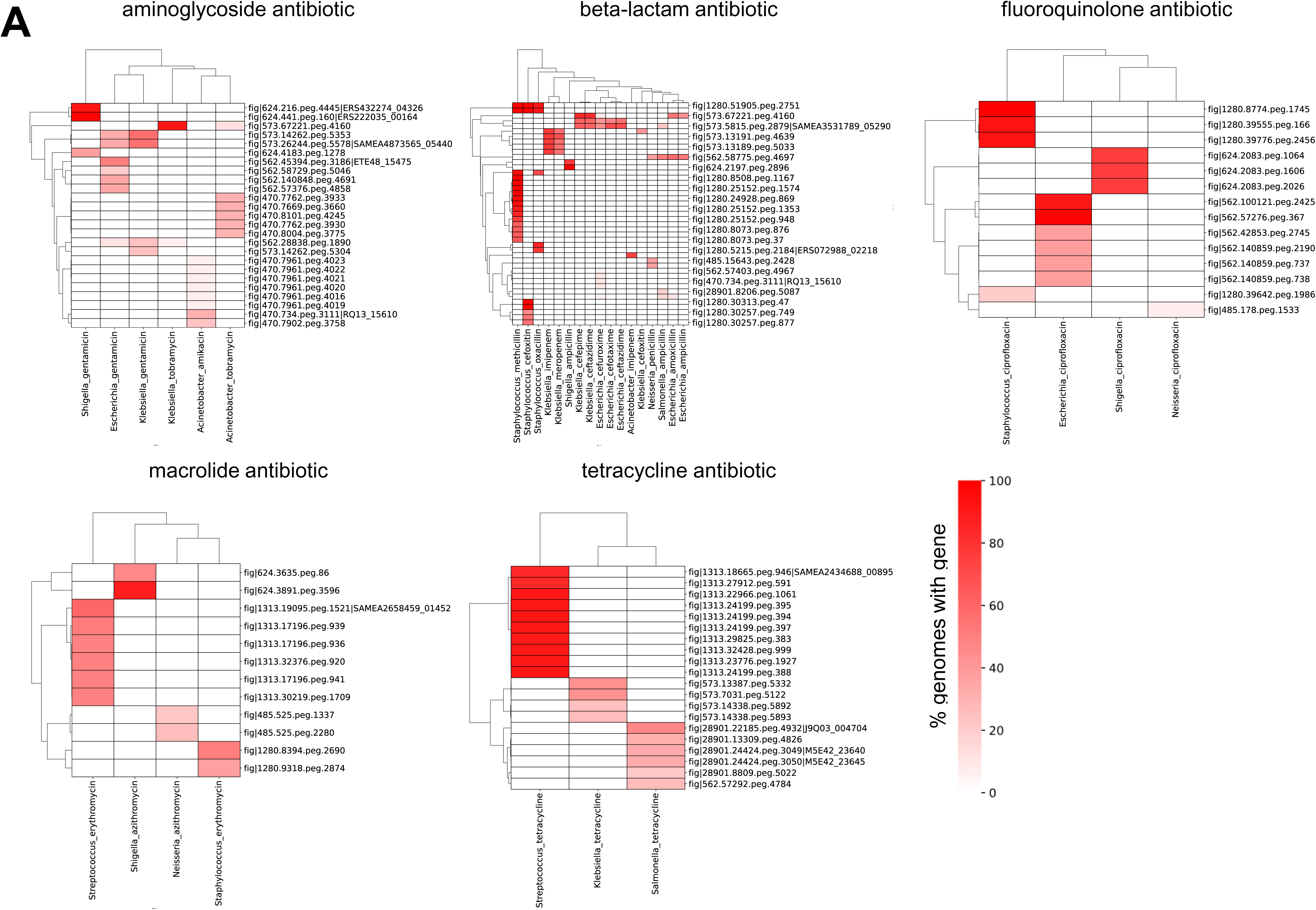
Clustermaps (by Euclidean distance) of novel AMR-associated gene abundance. Only genes present in ≥5% of genomes are shown.

**Supplemental Figure 1B.**
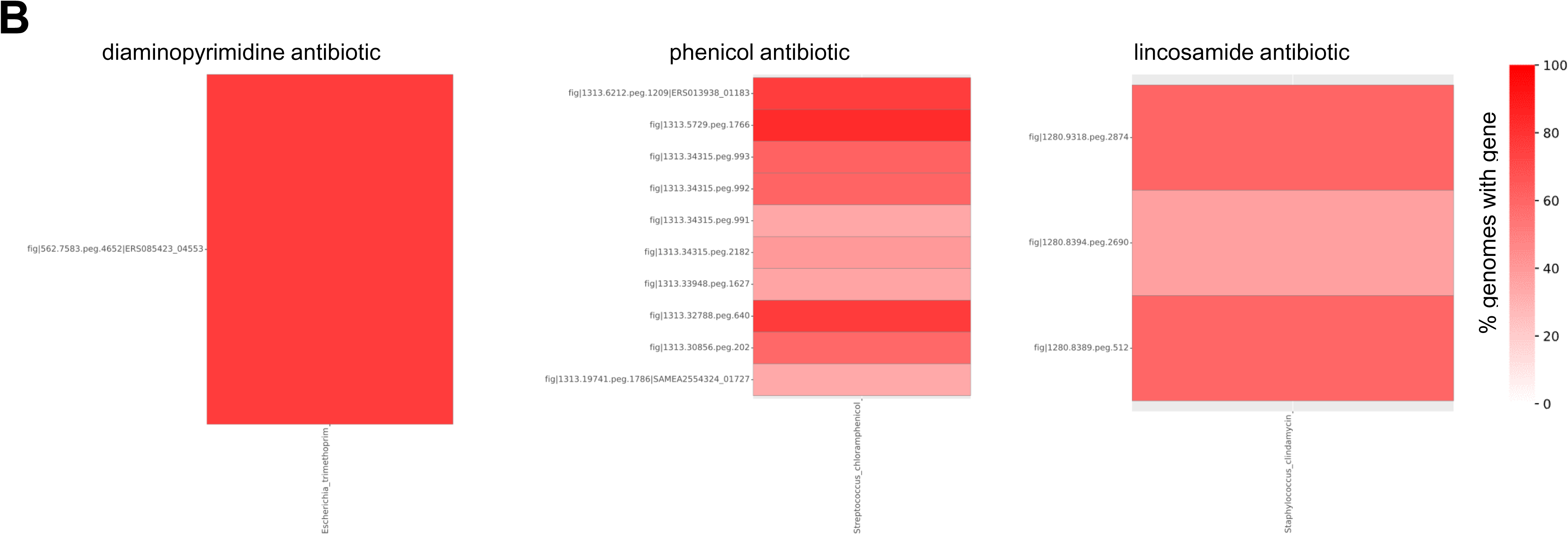
Heatmaps of novel AMR-associated gene abundance. Only genes present in ≥5% of genomes are shown.

**Supplemental Figure 2.**
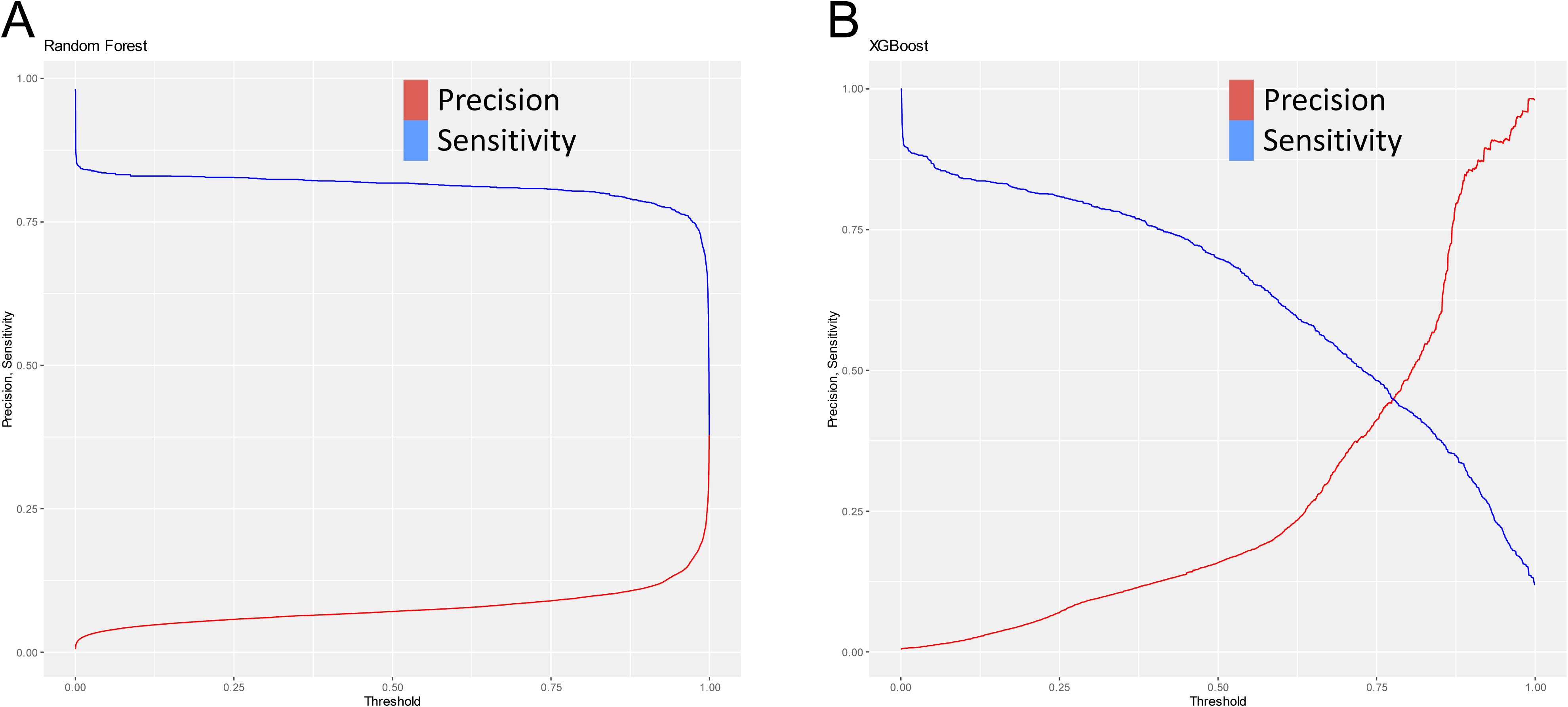
Comparison of precision versus sensitivity based on threshold for Random Forest (A) and XGBoost (B) classifier models.

**Supplemental Figure 3.**
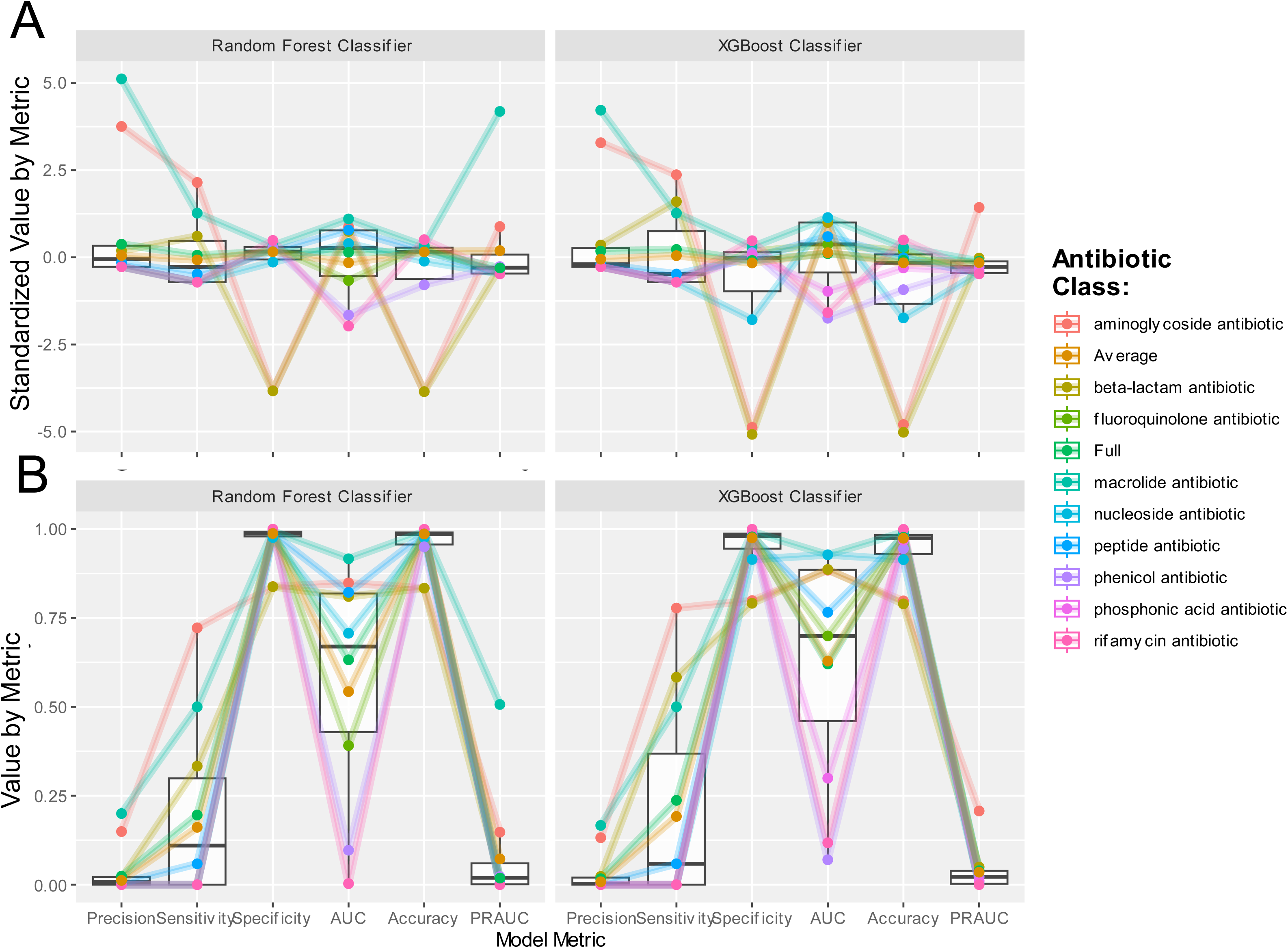
Comparison of standardized (A) and non-standardized (B) model metrics Random Forest (RF) and XGBoost (XG) classifiers across different classes in which both proteins and ligands not in training (Grp1).

**Supplemental Figure 4.**
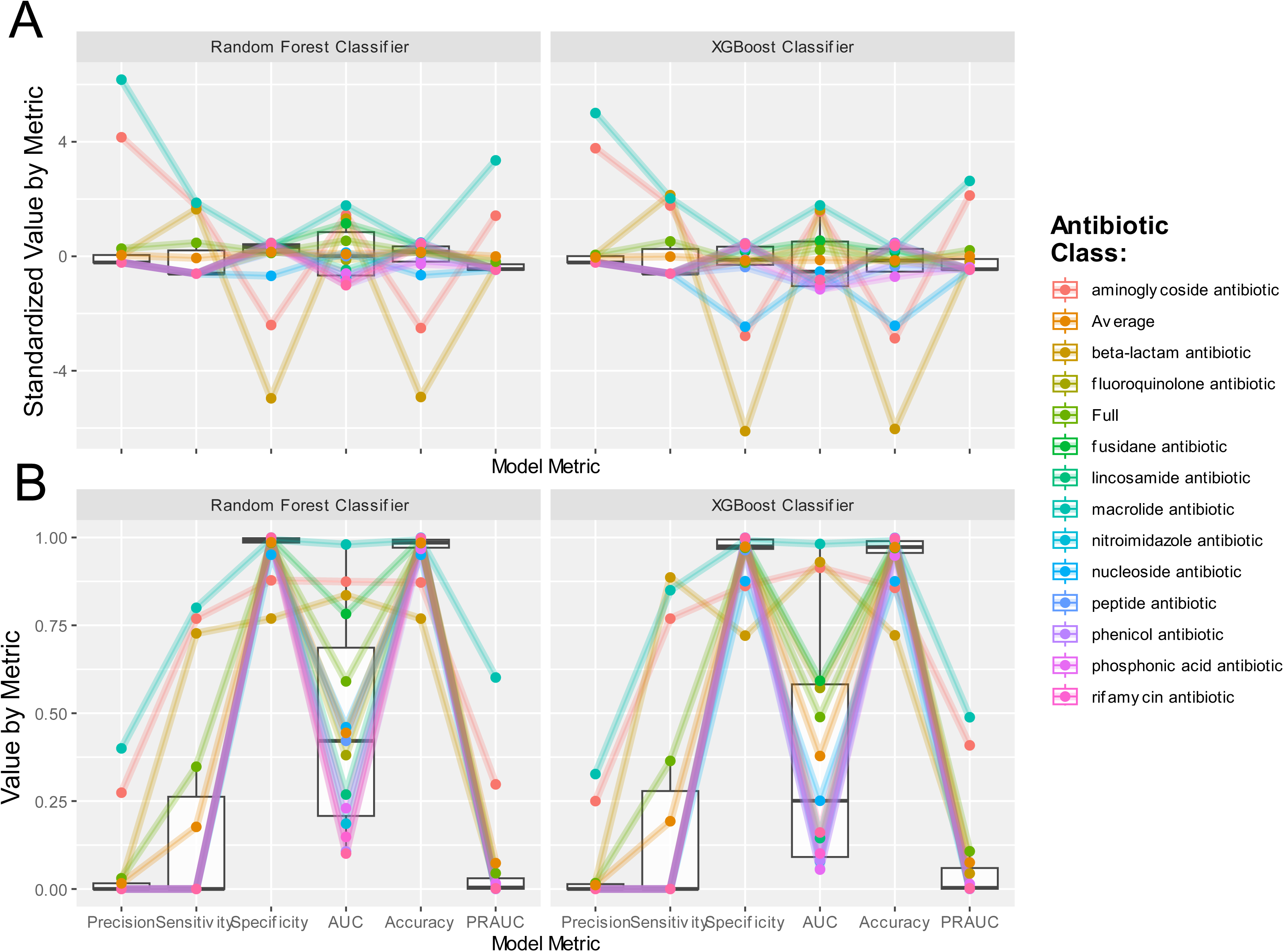
Comparison of standardized (A) and non-standardized (B) model metrics Random Forest (RF) and XGBoost (XG) classifiers across different classes in which proteins in training and ligands not in training (Grp2).

**Supplemental Figure 5.**
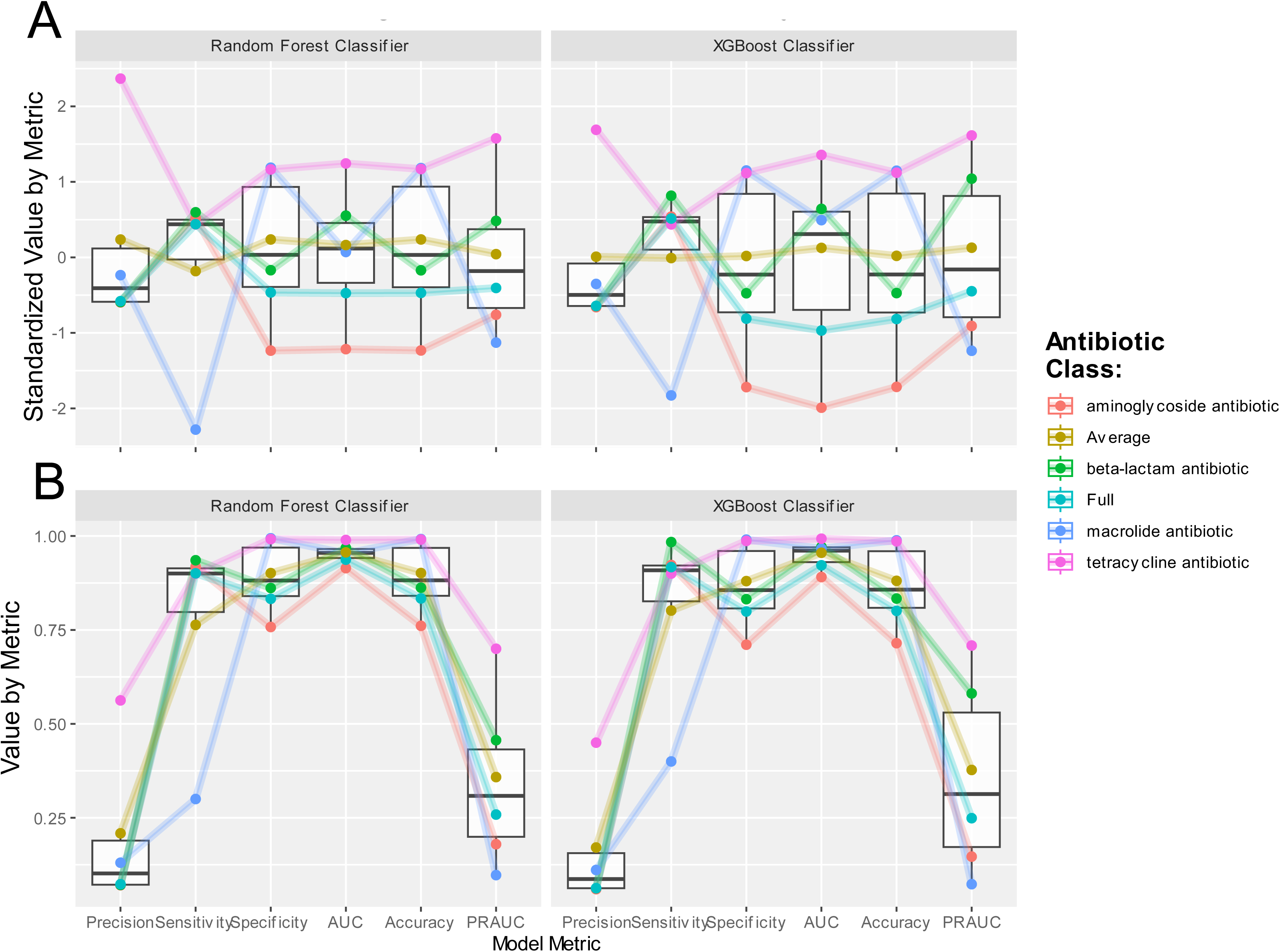
Comparison of standardized (A) and non-standardized (B) model metrics Random Forest (RF) and XGBoost (XG) classifiers across different classes in which ligands in training and proteins not in training (Grp3).

## References

1. Flynn, C. E. & Guarner, J. Emerging Antimicrobial Resistance. Mod. Pathol. 36, 100249 (2023).

2. Ho, C. S. et al. Antimicrobial resistance: a concise update. Lancet Microbe 6, (2025).

3. Darby, E. M. et al. Molecular mechanisms of antibiotic resistance revisited. Nat. Rev. Microbiol. 21, 280–295 (2023).

4. Hassall, J., Coxon, C., Patel, V. C., Goldenberg, S. D. & Sergaki, C. Limitations of current techniques in clinical antimicrobial resistance diagnosis: examples and future prospects. Npj Antimicrob. Resist. 2, 16 (2024).

5. Miller, W. R. & Arias, C. A. ESKAPE pathogens: antimicrobial resistance, epidemiology, clinical impact and therapeutics. Nat. Rev. Microbiol. 22, 598–616 (2024).

6. Waddington, C. et al. Exploiting genomics to mitigate the public health impact of antimicrobial resistance. Genome Med. 14, 15 (2022).

7. Sim, J., Kim, D., Kim, B., Choi, J. & Lee, J. Recent advances in AI-driven protein-ligand interaction predictions. Curr. Opin. Struct. Biol. 92, 103020 (2025).

8. Alcock, B. P. et al. CARD 2023: expanded curation, support for machine learning, and resistome prediction at the Comprehensive Antibiotic Resistance Database. Nucleic Acids Res. 51, D690–D699 (2023).

9. Olson, R. D. et al. Introducing the Bacterial and Viral Bioinformatics Resource Center (BV-BRC): a resource combining PATRIC, IRD and ViPR. Nucleic Acids Res. 51, D678–D689 (2023).

10. Jaillard, M. et al. A fast and agnostic method for bacterial genome-wide association studies: Bridging the gap between k-mers and genetic events. PLOS Genet. 14, e1007758 (2018).

11. Buchfink, B., Reuter, K. & Drost, H.-G. Sensitive protein alignments at tree-of-life scale using DIAMOND. Nat. Methods 18, 366–368 (2021).

12. Jones, P. et al. InterProScan 5: genome-scale protein function classification. Bioinformatics 30, 1236–1240 (2014).

13. Lin, Z. et al. Evolutionary-scale prediction of atomic-level protein structure with a language model. Science 379, 1123–1130 (2023).

14. Corso, G., et al. Deep Confident Steps to New Pockets: Strategies for Docking Generalization. (2024).

15. McNutt, A. T. et al. GNINA 1.0: molecular docking with deep learning. J. Cheminformatics 13, 43 (2021).

16. Papp, M. & Solymosi, N. Review and Comparison of Antimicrobial Resistance Gene Databases. Antibiotics 11, (2022).

17. Blake, K. S. et al. The tetracycline resistome is shaped by selection for specific resistance mechanisms by each antibiotic generation. Nat. Commun. 16, 1452 (2025).

18. Ebmeyer, S., Kristiansson, E. & Larsson, D. G. J. Unraveling the origins of mobile antibiotic resistance genes using random forest classification of large-scale genomic data. Environ. Int. 198, 109374 (2025).

19. Bush Karen & Bradford Patricia A. Epidemiology of β-Lactamase-Producing Pathogens. Clin. Microbiol. Rev. 33, 10.1128/cmr.00047-19 (2020).

20. Huang, Y. et al. Aminoglycoside-resistance gene signatures are predictive of aminoglycoside MICs for carbapenem-resistant Klebsiella pneumoniae. J. Antimicrob. Chemother. 77, 356–363 (2022).

21. Jiang, X. et al. Dissemination of antibiotic resistance genes from antibiotic producers to pathogens. Nat. Commun. 8, 15784 (2017).

22. Wencewicz, T. A. Crossroads of Antibiotic Resistance and Biosynthesis. Mol. Basis Antibiot. Action Resist. 431, 3370–3399 (2019).

23. Gibson, M., Nur-e-alam, M., Lipata, F., Oliveira, M. A. & Rohr, J. Characterization of Kinetics and Products of the Baeyer−Villiger Oxygenase MtmOIV, The Key Enzyme of the Biosynthetic Pathway toward the Natural Product Anticancer Drug Mithramycin from Streptomyces argillaceus. J. Am. Chem. Soc. 127, 17594–17595 (2005).

24. Tang, M.-C., Zou, Y., Watanabe, K., Walsh, C. T. & Tang, Y. Oxidative Cyclization in Natural Product Biosynthesis. Chem. Rev. 117, 5226–5333 (2017).

25. Gasparrini, A. J. et al. Tetracycline-inactivating enzymes from environmental, human commensal, and pathogenic bacteria cause broad-spectrum tetracycline resistance. *Commun*. Biol. 3, 241 (2020).

26. Park, J. et al. Plasticity, dynamics, and inhibition of emerging tetracycline resistance enzymes. Nat. Chem. Biol. 13, 730–736 (2017).

27. Maruyama, C. & Hamano, Y. The Biological Function of the Bacterial Isochorismatase-Like Hydrolase SttH. Biosci. Biotechnol. Biochem. 73, 2494–2500 (2009).

28. Vetting, M. W. et al. Mechanistic and Structural Analysis of Aminoglycoside N-Acetyltransferase AAC(6′)-Ib and Its Bifunctional, Fluoroquinolone-Active AAC(6′)-Ib-cr Variant. Biochemistry 47, 9825–9835 (2008).

29. Robicsek, A. et al. Fluoroquinolone-modifying enzyme: a new adaptation of a common aminoglycoside acetyltransferase. Nat. Med. 12, 83–88 (2006).

30. Zubyk Haley L. & Wright Gerard D. CrpP Is Not a Fluoroquinolone-Inactivating Enzyme. Antimicrob. Agents Chemother. 65, 10.1128/aac.00773-21 (2021).

31. Yagimoto, K., Hosoda, S., Sato, M. & Hamada, M. Prediction of antibiotic resistance mechanisms using a protein language model. Bioinformatics 40, btae550 (2024).

32. Sunuwar, J. & Azad, R. K. Identification of Novel Antimicrobial Resistance Genes Using Machine Learning, Homology Modeling, and Molecular Docking. Microorganisms 10, (2022).

33. Wong, F. et al. Benchmarking AlphaFold-enabled molecular docking predictions for antibiotic discovery. Mol. Syst. Biol. 18, e11081 (2022).

34. Villalba de la Peña, M. & Kronholm, I. Antimicrobial resistance in the wild: Insights from epigenetics. Evol. Appl. 17, e13707 (2024).

